# Conditioned pain modulation of pressure pain is associated with reduced activation in the parietal operculum and posterior insula

**DOI:** 10.1101/2023.08.23.554467

**Authors:** Karita E. Ojala, Ying Chu, Jürgen Finsterbusch, Christian Büchel

## Abstract

Conditioned pain modulation (CPM) is the phenomenon of pain inhibiting further pain. CPM is impaired in patients suffering from various chronic pain conditions and may resemble a pro-nociceptive profile. A possible neural mechanism for CPM is descending control from cortical regions via brainstem that modulates neural activity at the level of the dorsal horn of the spinal cord where noxious input first arrives. We studied the involvement of the descending pain modulatory pathway by simultaneous functional imaging of the brain, brainstem, rostral medulla, and cervical spinal cord during CPM with pressure cuff algometry in 42 healthy human participants. Subjective CPM as measured by decreased pain ratings developed over the experiment with application of repeated short test stimuli for long conditioning pain compared with non-painful conditioning. Test stimuli paired with painful conditioning were associated with decreased BOLD fMRI activity in the parietal operculum (secondary somatosensory cortex) and posterior insula, as well as increased responses in bilateral primary somatosensory cortices. Subjective CPM reflected in pain ratings was associated with decreased test stimulus responses in somatosensory and pain-responsive brain regions such as the anterior and posterior insula and the parietal operculum. Conditioning pain and test pain were associated with increased neural activity in stimulus-ipsilateral spinal cord dorsal horn at segment C6 and C4, respectively. No evidence for neural correlates of CPM or associations with subjective CPM was observed in the spinal cord. Functional connectivity between the brain and spinal cord was altered during both conditioning and test pain but did not depend on conditioning intensity. Taken together, we observed neural activity associated with both negative (CPM) and positive modulation of pain in different somatosensory and pain-associated brain regions, and an association of these and other neural responses with subjective CPM. While we found that painful stimuli altered the connectivity along the descending pain modulatory pathway, we did not find evidence for a modulation by conditioning intensity nor neural correlates of CPM in the spinal cord.

## Introduction

Pain is a central control signal that allows an organism to avoid bodily harm. However, it is also beneficial for the organism to be able to modulate how intensely pain is felt, depending on the situation. For example, persistent unavoidable pain that is not life-threatening may be inhibited as its utility as a learning signal decreases especially in the presence of more threatening stimuli [1]. Descending pathways from the cerebral cortex to the spinal cord can modulate pain by top-down inhibition of noxious input, involving various neurotransmitters such as endogenous opioids, endocannabinoids, and monoamines [2–4]. Cortical regions involved in pain modulation include anterior cingulate cortex (ACC), ventromedial prefrontal cortex (VMPFC) and insula, which take part in pain valuation, learning, prediction, and expectations [1,4]. Additionally, dorsolateral prefrontal cortex (DLPFC) may take part in pain suppression and maintenance of pain inhibition [5]. These cortical regions send descending signals via intermediate nodes in the brainstem and medulla (periaqueductal grey, PAG; locus coeruleus; rostroventral medulla, RVM) to modulate pain transmission in the dorsal horn of the spinal cord where noxious input first enters the central nervous system [1,2,4]. In addition, dorsal horn signals are also modulated within the spinal cord, both across segments of the spinal cord via propriospinal tracts [6] as well as locally within segments [7].

A particular form of endogenous pain modulation is the phenomenon of “pain inhibits pain”, where pain sensation is inhibited by preceding or simultaneous painful stimuli in a distal location such as another limb. In rodents and primates, the underlying neural mechanism is termed diffuse noxious inhibitory controls (DNIC), where the responses of second-order wide-dynamic range neurons in the spinal cord dorsal horn are inhibited after noxious input outside their receptive fields [8,9]. DNIC has a supraspinal origin [10] and is mediated by brainstem noradrenergic circuits [11,12] and the subnucleus reticularis dorsalis (SRD) in caudal medulla [6,9,13] but also by descending endogenous opioid pathways [14].

In humans, experimental paradigms where one painful stimulus inhibits another are referred to as conditioned pain modulation (CPM) and it has been suggested that the individual CPM capacity may reflect an individual marker of pain processing (i.e., “pro-nociceptive profile” with impaired pain inhibition as measured with CPM), which in turn may be associated with the risk of developing chronic pain [15]. Less efficient CPM has been shown to be predictive of development of post-operative chronic pain, and patients suffering from some forms of chronic pain, namely idiopathic pain syndromes such as fibromyalgia, migraine, and osteoarthritis, have shown to exhibit decreased CPM in some studies [15,16]. However, it may be the case that chronic pain also changes an individual’s capacity for CPM [15], and the evidence of CPM’s association with features of chronic pain such as intensity, disability and duration is mixed [17]. While CPM’s role as a potential biomarker for chronic pain needs further investigation, it is a widely applied method for investigating individual pain modulation capacity in healthy individuals and patients.

The precise neural mechanism of human CPM, and to what extent it may be similar to DNIC in animal models, is not known [3]. While CPM may involve a DNIC-like mechanism in humans, paradigms in awake humans likely include a wide range of factors that can influence pain perception, such as attention and expectations [18]. Noxious pressure conditioning leading to CPM in humans has been shown to cause DNIC in rats in a comparable paradigm [19]. Only a few studies exist that have investigated the neural basis of CPM in humans [20] and only some of them have measured functional neural activity during CPM [21–26]. In half or more of these studies, CPM was associated with decreased BOLD signals in the thalamus and anterior insula, while fewer studies found decreased activity in mid-cingulate cortex, posterior cingulate cortex, posterior insula, secondary somatosensory cortex, and inferior frontal gyrus [20]. Increased activity has been mostly observed in superior temporal gyrus and primary somatosensory cortex but also in superior frontal gyrus [20]. Some studies have also explored brainstem activity related to CPM and found association of CPM magnitude with BOLD responses in brainstem and medullar nuclei including parabrachial nucleus and SRD, and BOLD activity decreases in the parabrachial nucleus and PAG that correlated with behavioural CPM response [24,27]. However, there is large variability in the experimental paradigms, and sample sizes were often small [20,28]. Moreover, studies including a control condition for the conditioning stimulus, and/or a direct comparison of neural activity for test stimuli with conditioning to test stimuli with no (painful) conditioning would be important to precisely disentangle CPM effects from pain responses alone [20]. Finally, neural correlates of CPM have not been previously investigated in the entire central nervous system also covering the spinal cord, which would allow integrating information across the entire descending pain modulatory pathway.

Combined functional magnetic resonance imaging (fMRI) of the brain and spinal cord allows near-simultaneous acquisition of functional images at several different levels of the central nervous system, making it a is a powerful tool for investigating neural function and connectivity across the descending pain modulatory pathway [29]. Here, we used cortico-spinal fMRI to acquire BOLD signals from nearly the entire brain, brainstem, caudal medulla, and relevant segments of the cervical spinal cord while healthy male and female participants underwent blocks of tonic painful or non-painful pressure on the left arm while also receiving painful phasic pressure on the right arm (Figure 1). The pressure pain was applied with a cuff algometer that can induce deep tissue pain in addition to superficial pain. Ischemic pressure pain applied with a cuff algometer has been shown to reliably induce CPM in humans [30–32]. The simultaneous application of tonic and phasic stimuli was motivated by the requirement for DNIC mechanism that noxious stimuli are applied in a concurrent and competing manner [33] and allowed us to acquire trial-wise pain ratings as well as BOLD responses over multiple stimulus repetitions. We investigated whether distinct neural correlates of test and conditioning stimuli as well as correlates of CPM could be observed in the brain, but also in the brainstem, medulla and spinal cord. Finally, we explored interactions between these levels of the descending pain modulatory pathway.

**Figure 1.**
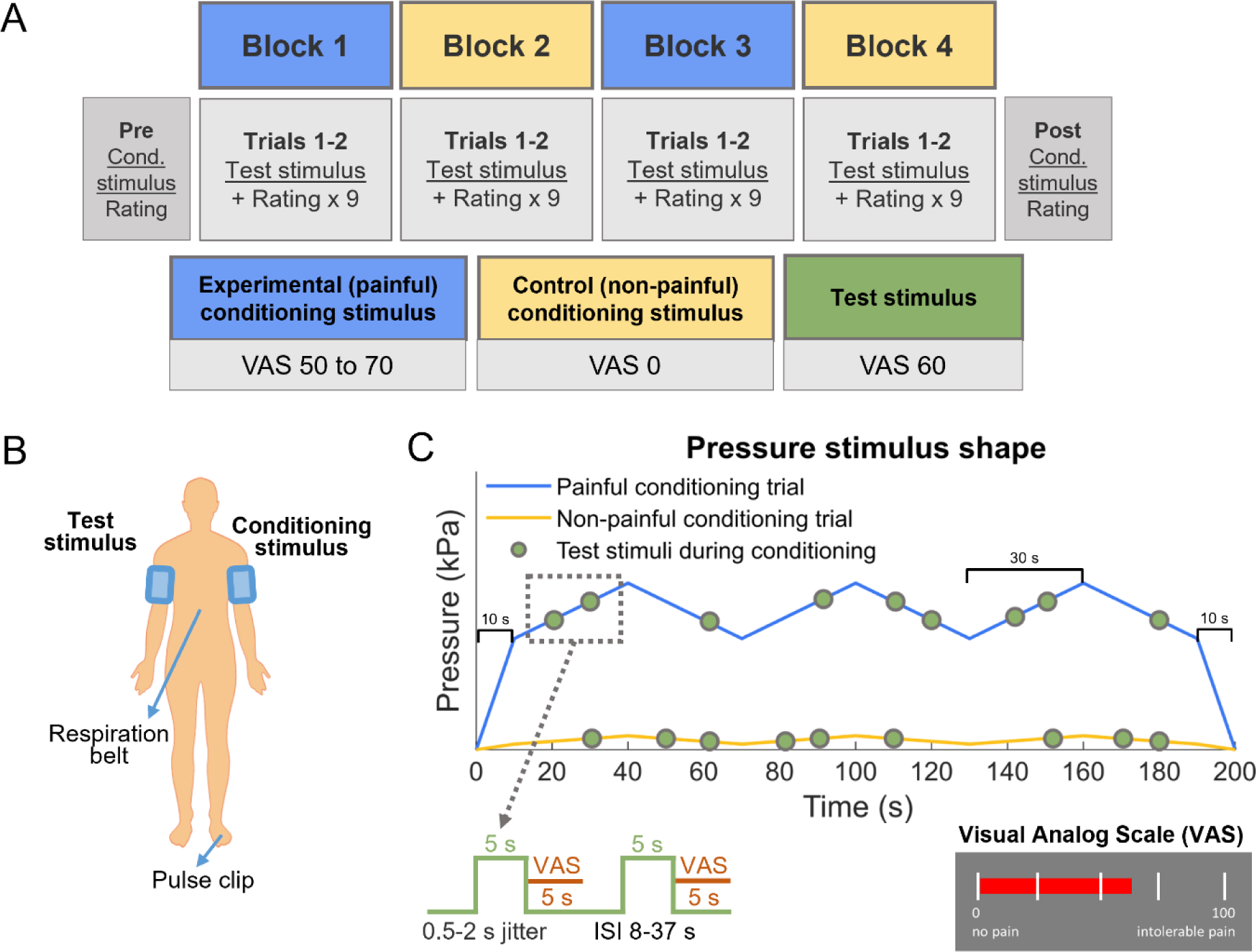
Experimental design, pressure cuff and psychophysiological measurement placement, and pressure stimulus shape during a trial. (**A**) The experiment consisted of a pre-experiment continuous pain rating of the painful conditioning stimulus, four experimental blocks with concurrent short test stimuli and long-lasting conditioning stimuli, and a post-experiment continuous pain rating of the painful conditioning stimulus. Two of the experimental blocks contained painful conditioning stimulation and two non-painful conditioning stimulation. The order of the blocks was counter-balanced across participants. Each block contained two trials of conditioning stimulus and concurrent repeating painful short test stimuli, followed by a VAS pain rating period. (**B**) Conditioning stimuli were applied on the right arm and test stimuli on the left arm via automatized pressure cuff algometry. Respiration was measured with a belt around the chest and pulse via a photoplethysmograph on the left big toe during the entire experiment. (**C**) Painful conditioning stimuli fluctuated slightly, completing three cycles within one stimulus with a total duration of 200 s. The non-painful conditioning stimulus also fluctuated slightly with very low pressure output. Three test stimuli of 5 s duration were applied on each of the three cycles of each conditioning stimulus, for a total of 9 test stimuli per conditioning stimulus and 18 test stimuli per block, for 72 total test stimuli over the entire experiment. The pain rating was done with the right hand within 5 s.

## Results

### Effect of combined conditioning and test pressure on pain ratings

Firstly, we were interested to see whether conditioning with tonic pressure pain, in comparison with non-painful tonic pressure, led to decreased pain ratings to phasic test pressure stimuli – potentially reflecting behavioural CPM. Since the experiment consisted of 72 test stimuli over 4 blocks, we also expected that time or the number of stimulus presentations (trials) may also be important (e.g., habituation). To investigate these effects, we first looked at the mean test stimulus pain ratings during painful vs. non-painful conditioning stimulus. Without accounting for trials, there was no mean difference in test pain ratings between the conditioning stimulus conditions (0.9 VAS, SD = 9.3, *t*(41) = 0.63, *p* = 0.27). However, we observed that test stimulus pain ratings were lower during painful conditioning than non-painful conditioning when comparing late to early trials (Figure 1A). Linear mixed effects model confirmed this observation. In addition to main effects of conditioning stimulus intensity (non-painful > painful), F(1,2902) = 4.22, p = 0.04 and number of test stimuli, F(1,2902) = 7.84, p = 0.005, there was a significant interaction indicating that painfulness of the test stimuli decreased more steeply for painful than non-painful conditioning stimuli during the experiment (Figure 2B, interaction effect *F*(1,2902) = 12.56, *p* = 0.0004). In absolute terms, test pain decreased on average by 14.4 VAS units from the first five painful conditioning test ratings to the last five, and 6.3 VAS units from first five non-painful conditioning test rating to the last five, giving a 23.4 % vs. 11.6 % decrease comparing to the mean of first five ratings (Figure 2A). That is, test pain ratings decreased on average around 11.8 % points more during painful than non-painful conditioning.

**Figure 2.**
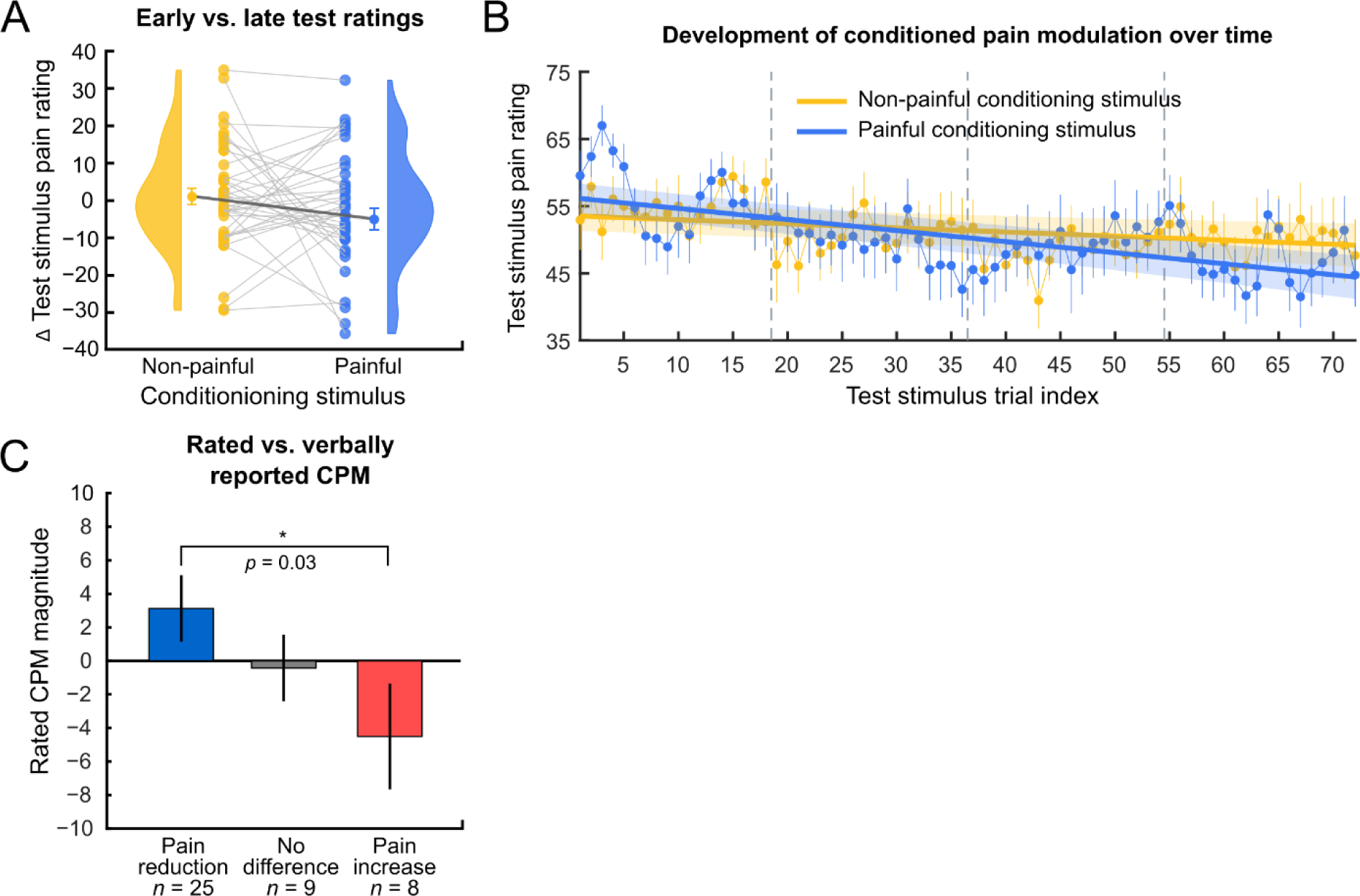
Pain ratings for test stimuli and post-experiment verbal report of conditioned pain modulation (CPM). **(A)** The change in participant-wise means from early (first five) to late (last five) test stimulus pain ratings during non-painful and painful tonic conditioning stimuli. While pain ratings to test stimuli decreased more from the start to the end of the experiment for painful than non-painful tonic conditioning stimulus, there was large individual variation in effect magnitude and direction. **(B)** Development of CPM over all 72 test stimulus trials in the experiment. Solid thick lines show the model fit separately for painful and non-painful conditioning stimuli over test stimuli. The significant decrease in pain ratings over time was more pronounced for painful than for non-painful conditioning stimulus (interaction effect *p* < 0.001). (**C**) Comparison of CPM effect as measured by test stimulus pain ratings (non-painful – painful conditioning stimulus condition) and post-experiment verbal report for overall difference in pain in the test stimulus arm during non-painful vs. painful pressure on the conditioning stimulus arm. Sixty percent of participants (25 out of 42) reported hypoalgesia, and CPM (reduced pain) was significantly higher in those participants who reported hypoalgesia than in those who reported hyperalgesia. All error bars and areas show the standard error of the mean.

We were also interested whether trial-wise test pain ratings and post-experiment verbal report would be consistent with each other. Out of 42 participants, 25 reported hypoalgesia (pain on the right test stimulus arm was less with painful conditioning stimulus on the left arm), 9 reported not feeling any difference (pain was the same regardless of which conditioning stimulus was on), and 8 reported hyperalgesia (pain was worse with painful conditioning stimulus on). Trial-wise rated CPM magnitude (test pain ratings during non-painful – painful conditioning) was significantly higher in the group of participants who reported hypoalgesia than in those who reported hyperalgesia (Figure 2C, t(13) = 2.05, p = 0.031, Cohen’s d = 0.79).

Finally, there was no significant correlations of average rated CPM with any of the following background variables: sex, age, body mass index, nor with any of the following features of the experimental paradigm: arm calibration order, first block conditioning stimulus intensity (painful or non-painful), last block conditioning stimulus intensity (Table S1). Verbal CPM was only positively correlated with test stimulus pressure, *p* = 0.013, and negatively correlated with test stimulus calibration regression slope, *p* = 0.018 (Table S1; not corrected for multiple comparisons).

### Brain activity related to tonic conditioning pain

Since we used a novel type of fluctuating tonic pressure stimulus, we first confirmed that this stimulus was associated with BOLD fMRI responses in brain regions commonly activated during pain. Whole-brain analysis with Threshold-Free Cluster Enhancement (TFCE), voxel-wise family-wise error (FWE)-corrected at p < 0.025 for comparisons in both contrast directions, showed significantly increased activity during painful than non-painful conditioning stimulus (‘Painful > non-painful conditioning stimulus’ contrast; Figure 3A) in bilateral primary somatosensory (S1) and motor (M1) cortices, bilateral supplementary motor area (SMA) and anterior paracentral lobule, left superior parietal lobule and precuneus, right anterior insula, posterior mid-cingulate cortex (MCC), anterior ventromedial prefrontal cortex (vmPFC), as well as left superior-middle frontal gyrus/dorsolateral prefrontal cortex (dlPFC). Increased BOLD responses for the non-painful than for painful conditioning stimulus was observed in the cerebellum. Our pre-registered, a priori anatomical regions-of-interest (ROIs) included brain regions previously associated with pain processing and modulation, namely, secondary somatosensory cortex (S2)/parietal operculum, anterior and posterior insula, rACC, vmPFC, dlPFC, PAG in the brainstem and RVM. We extracted BOLD signal parameter estimates (betas) from each of these ROIs, compared mean estimates in a within-subject manner, and found that BOLD signal in the dlPFC, t(41) = 2.99, p = 0.0023, Cohen’s d = 0.46, and vmPFC, t(41) = 3.53, p = 0.0005, Cohen’s d = 0.54 (p < 0.0045 considered significant Bonferroni-corrected for the number of ROIs), was significantly higher during painful than non-painful conditioning stimuli (Figure 3B). There were no significant differences in other brain ROIs.

**Figure 3.**
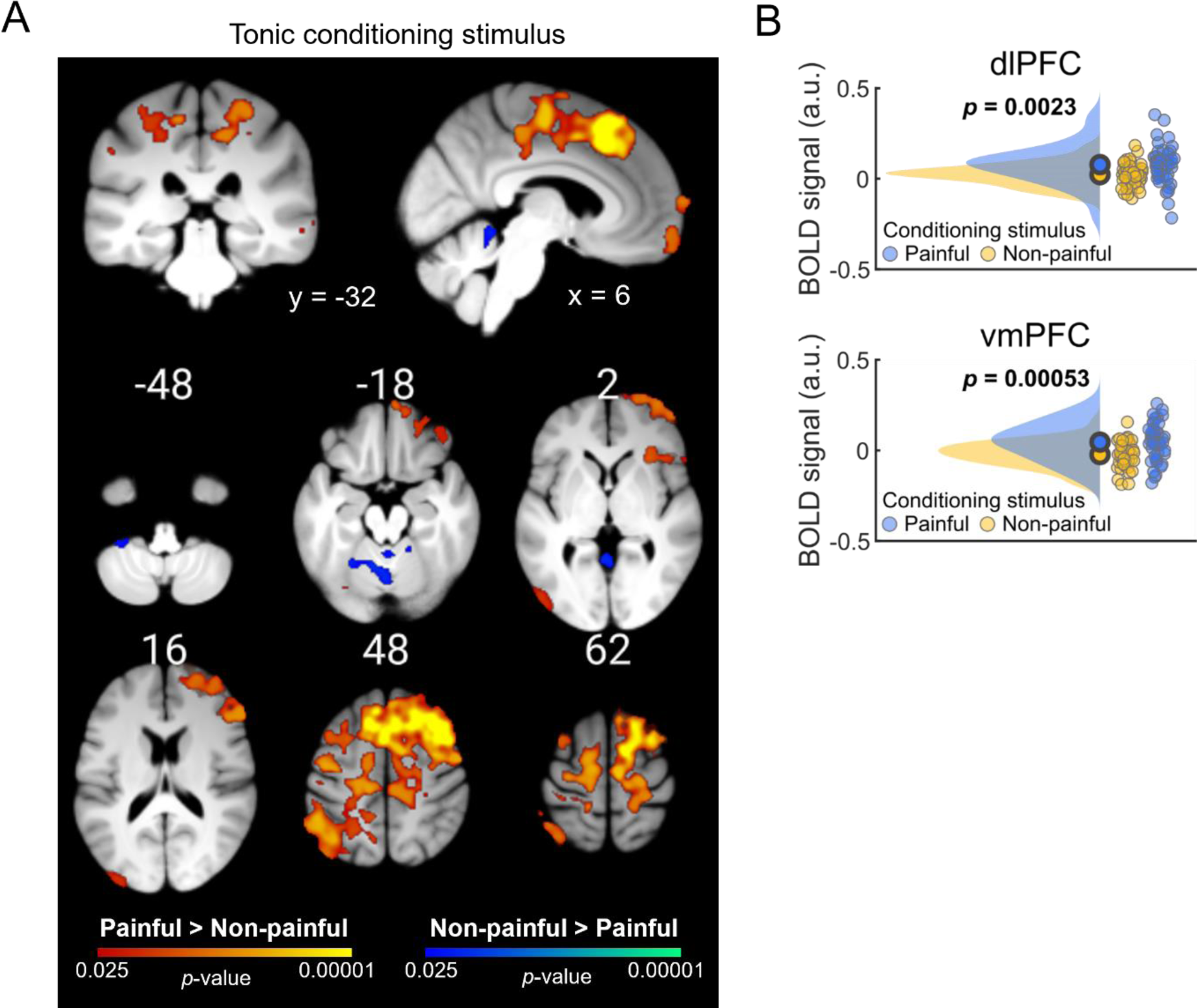
Brain BOLD fMRI activity associated with tonic conditioning stimulus on the left arm. **(A)** Whole-brain analysis results with Threshold-Free Cluster-Enhancement (TFCE), family-wise error (FWE)-corrected at *p* < 0.025 (corrected for two-tailed one-sample t-test). Significantly higher activity for painful than non-painful conditioning stimulus was observed in a wide range of sensorimotor and pain-associated regions, such as bilateral primary somatosensory (S1) and motor (M1) cortices, bilateral supplementary motor area (SMA) and anterior paracentral lobule, left superior parietal lobule and precuneus, right anterior insula, posterior mid-cingulate cortex (MCC), anterior ventromedial prefrontal cortex (vmPFC), as well as left superior-middle frontal gyrus/dorsolateral prefrontal cortex (dlPFC). Activity higher for the non-painful stimulus was observed only in the cerebellum. **(B)** BOLD signal parameter estimates extracted from a priori anatomical regions-of-interest. Significant differences between parameter estimates for painful and non-painful conditioning stimuli were only seen in vmPFC and dlPFC (bilateral ROIs) where activity was increased for the painful stimulus relative to the non-painful stimulus (one-tailed paired t-test, *p* < 0.0045 considered significant when correcting for 11 ROIs). a.u. = arbitrary units.

The fluctuation of the conditioning stimulus was included as we had an exploratory hypothesis that we may be able to observe changes in the BOLD signal corresponding to the changes in the tonic pressure. However, this was not the case, and we do not report results on the tonic pressure fluctuation in any further analyses.

### Spinal cord activity related to tonic conditioning pain

Next, we investigated whether the conditioning stimulus was associated with changes in BOLD activity in the spinal cord. We conducted a small volume correction for the left side (i.e. ipsilateral to the side of the tonic stimulus) of the dorsal horn across segments C4, C5 and C6 with TFCE at *p* < 0.05 FWE-corrected (one-tailed paired t-tests; we were not interested in the other direction in this case). Our data show that the BOLD signal was significantly increased in right side of segment C5 during non-painful conditioning stimulus (‘Non-painful conditioning stimulus onset > baseline’ contrast; Figure 4A top row; around coordinates x = -4, y = -45, z = -153). The lower segment C6 also showed increased BOLD responses during the painful conditioning stimulus (‘Painful conditioning stimulus onset > baseline contrast; Figure 4A middle row; around coordinates, x = 0, y = -48, z = -173). In the same area, we also show higher responses to painful than non-painful conditioning stimuli, but these were non-significant at *p* < 0.05 FWE-corrected (‘Painful > non-painful conditioning stimulus onset’ contrast; Figure 4A bottom row; around coordinates x = -1, y = -48, z = -174; highest *p*-value around 0.108 while with traditional SPM approach with small volume correction we observed a peak with *p* = 0.028 voxel-wise FWE-corrected). We also extracted BOLD signal parameter estimates from the left dorsal horn segments C4, C5 and C6 and show a trend-level result of painful conditioning stimulus being associated with higher activity than non-painful conditioning stimulus in left dorsal horn at C6, *t*(41) = 2.15, *p* = 0.019, Cohen’s *d* = 0.33 (Figure 4B; *p* < 0.0167 considered significant corrected for the three spinal cord segments tested). While it was not part of our hypothesis, surprisingly we also observed a similar result for right side dorsal horn activation at C6, *t*(41) = 2.19, *p* = 0.017, *d* = 0.34). A further repeated-measures ANOVA also showed a trend-level main effect of conditioning stimulus condition, *F*(1,41) = 5.18, *p* = 0.028, 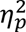 0.11, and no difference between sides of the spinal cord, F(1,41) = 1.37, *p* = 0.25, no interaction, *p* = 0.98, against our initial hypothesis of increased responses to painful over non-painful conditioning only in the left dorsal horn, ipsilateral to the conditioning stimulus arm (Figure 4C).

**Figure 4.**
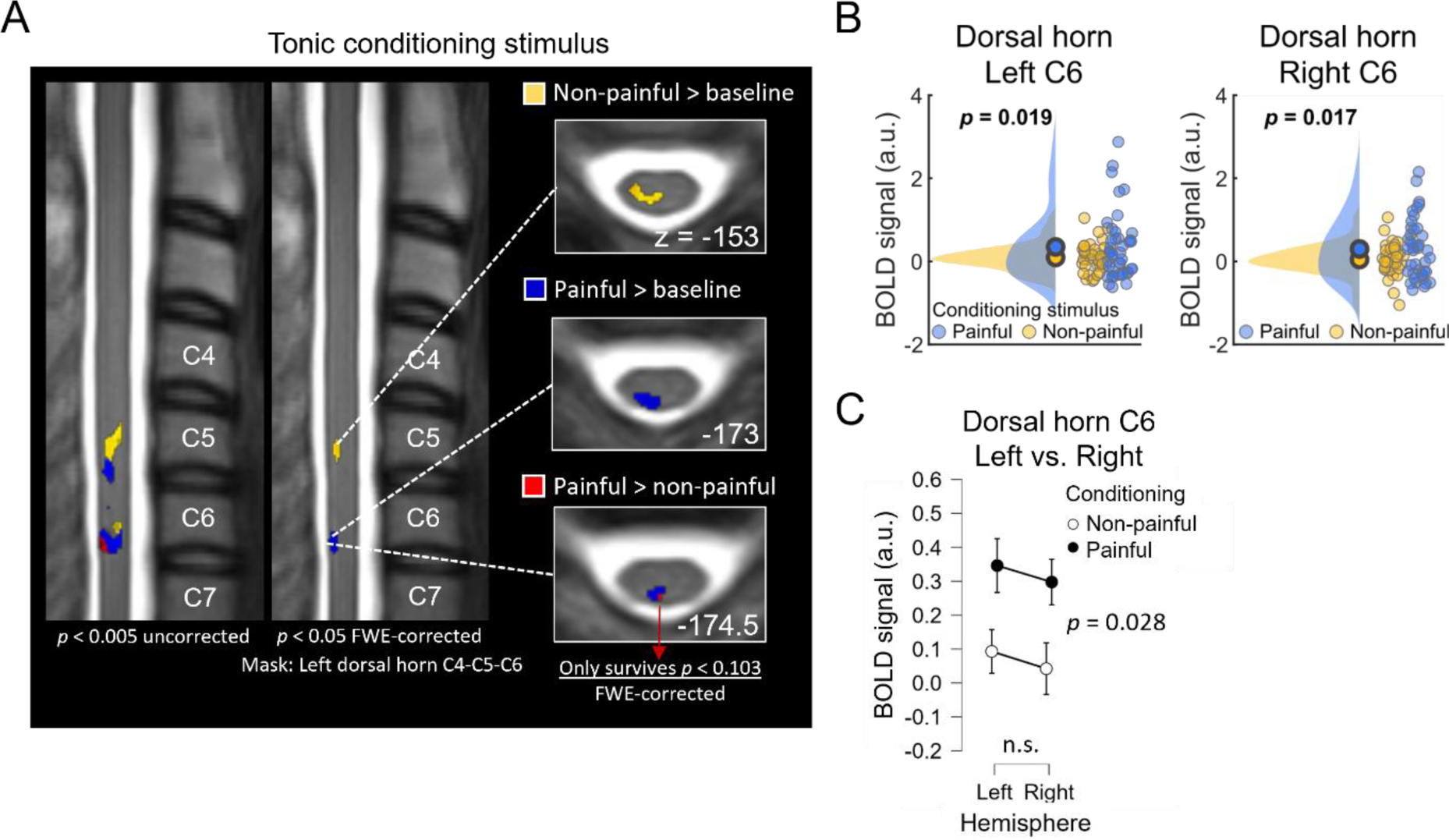
Spinal BOLD fMRI activity associated with tonic conditioning stimulus on the left arm. (**A**) Spinal cord analysis with TFCE, FWE-corrected at *p* < 0.05 (one-tailed one-sample t-test) within a priori hypothesized left dorsal horn segments C4, C5, and C6. Non-painful conditioning stimuli were associated with activity in left dorsal horn at segment C5, and painful conditioning stimuli at C6. In left dorsal horn at C6, painful conditioning stimuli were associated with increased activity compared to non-painful stimuli, but this comparison was not significant using FWE-correction at *p* < 0.05 as the smallest *p*-value for this contrast was 0.108. (**B**) BOLD signal parameter estimates extracted from a priori anatomical regions-of-interest in the dorsal horn. Increased signal for painful over non-painful conditioning stimulus was seen in left dorsal horn at segment C6 at trend-level (one-sided paired t-tests; *p* < 0.0167 considered significant when correcting for three spinal cord segments). However, when we explored our data beyond our initial hypothesis, we also found a very similar trend-level result on the right side of dorsal horn at C6. (**C**) In a repeated-measures ANOVA including factors tonic conditioning stimulus intensity and side of the spinal cord, there was a trend-level difference between painful and non-painful conditioning stimuli but no difference between sides of the dorsal horn.

### Brain activity related to phasic pressure pain and its combination with painful tonic pressure

Before analysing neural responses associated with CPM, we also wanted to confirm BOLD responses to the test stimuli on the right arm. In a whole-brain analysis with TFCE, we found increased BOLD activity in various regions associated with sensorimotor and pain processing, as well as decreased activity in visual and default mode network regions in response to the test stimuli in general, regardless of conditioning stimulus intensity (‘Test stimulus onset during painful & non-painful conditioning stimuli > baseline’ contrast; Figure 5).

**Figure 5.**
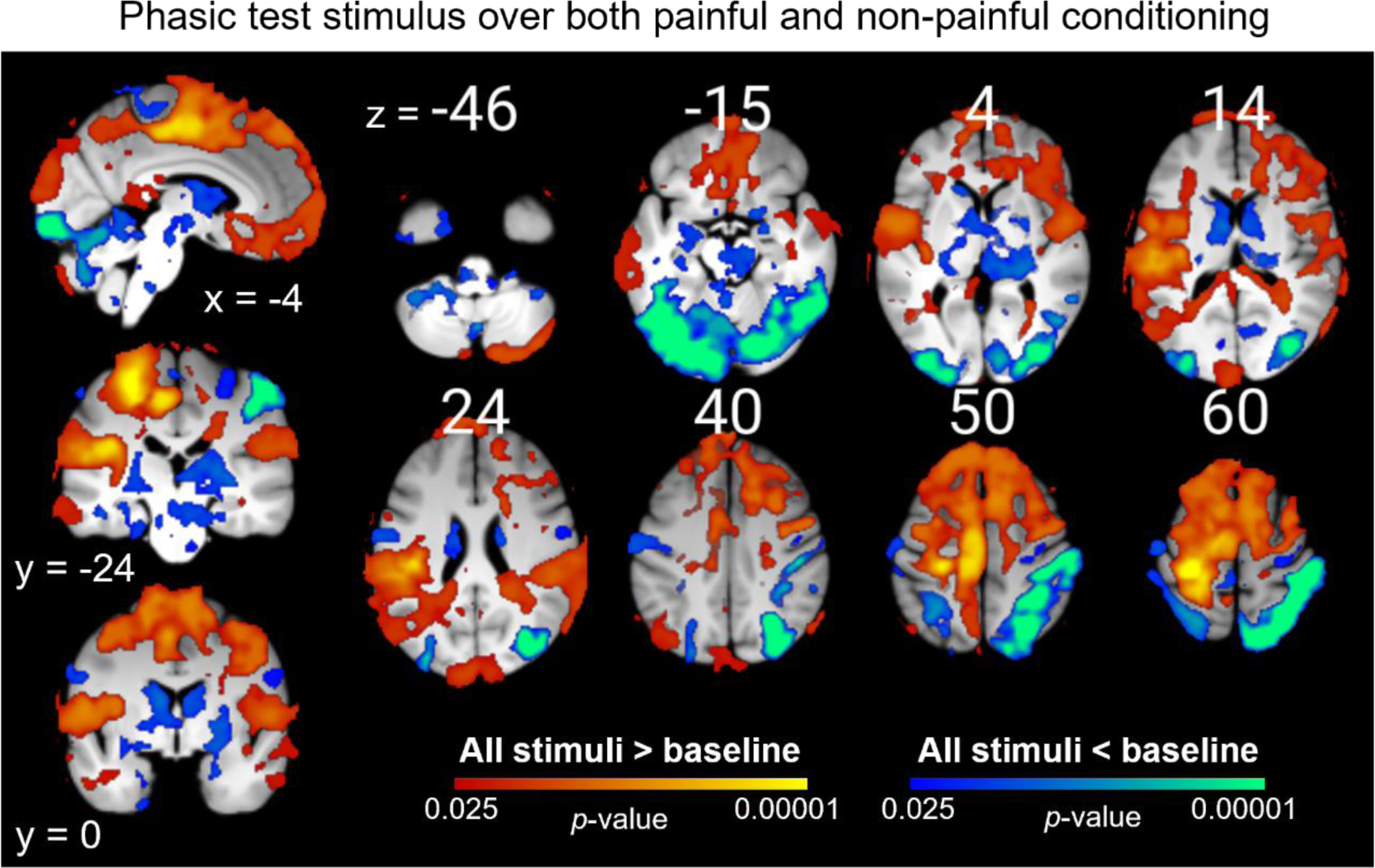
Brain BOLD fMRI activity associated with phasic test stimulus on the right arm regardless of conditioning stimulus. Whole-brain analysis results with TFCE, FWE-corrected at *p* < 0.025 (corrected for two-tailed one-sample t-test). Widespread activity with increases in sensorimotor and pain-associated networks and decreases in visual and default mode network regions was found in response to the test stimuli in general, regardless of conditioning stimulus intensity.

Central to our CPM hypothesis, we expected to observe either decreased (CPM, negative modulation) or increased (positive modulation) BOLD signals to phasic test pain during painful vs. non-painful tonic conditioning pressure in regions involved with pain processing and modulation, including left and right anterior and posterior insula, left and right parietal operculum (secondary somatosensory cortex, SII), and bilateral rACC, vmPFC, dlPFC, PAG and RVM. In a whole-brain analysis, we found that neural activity in regions encompassing bilateral S1 and superior parietal lobule, left M1, and left anterior paracentral lobule was higher for test stimuli during painful than non-painful conditioning stimuli (positive modulation, i.e., more signal for phasic stimulus in the presence of a painful tonic stimulus; Figure 6A). On the other hand, we found lower BOLD responses for test stimuli during painful in comparison with non-painful conditioning stimulus in right parietal operculum (CPM-like effect, or hypoalgesia due to conditioning; Figure 6A). Comparison of BOLD signal estimates extracted from our anatomical a priori brain ROIs for the test stimuli showed significantly lower responses during painful than non-painful conditioning stimuli in right parietal operculum, *t*(41) = −3.31, *p* = 0.002, Cohen’s *d* = 0.51, and posterior insula, *t*(41) = −4.95, *p* = 0.0001, *d* = 0.76 (significant at p < 0.0045 corrected for 11 ROIs; Figure 6B), while there were no significant differences for the left hemisphere, *p* = 0.20 for parietal operculum and *p* = 0.035 for posterior insula. This was interesting since our expectation was to observe a CPM effect (test stimuli during non-painful > painful conditioning) in the test stimulus-contralateral pain processing regions in the brain. Repeated-measures ANOVA revealed a difference for test stimulus activity between the hemispheres, *F*(1,41) = 78.00, *p* = 5e^-11^, 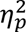 0.66, as well as between painful and non-painful conditioning stimuli, *F*(1,41) = 5.20, *p* = 0.028, 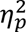 = 0.11 (only trend-level when corrected for number of ROIs) for parietal operculum, as well as for posterior insula, hemisphere effect *F*(1,41) = 11.8, *p* = 0.001, 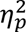 = 0.22, and conditioning stimulus effect *F*(1,41) = 14.5, *p* = 0.0005, 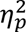 = 0.26, but no significant interaction of hemisphere and conditioning stimulus condition (parietal operculum *p* = 0.29, posterior insula *p* = 0.34). Finally, we observed no significant BOLD activity associated with test stimuli over time, that is, dependent on number of test stimulus presentations (‘test trial index’ related contrasts) in contrast to our finding in subjective trial-wise test pain ratings.

**Figure 6.**
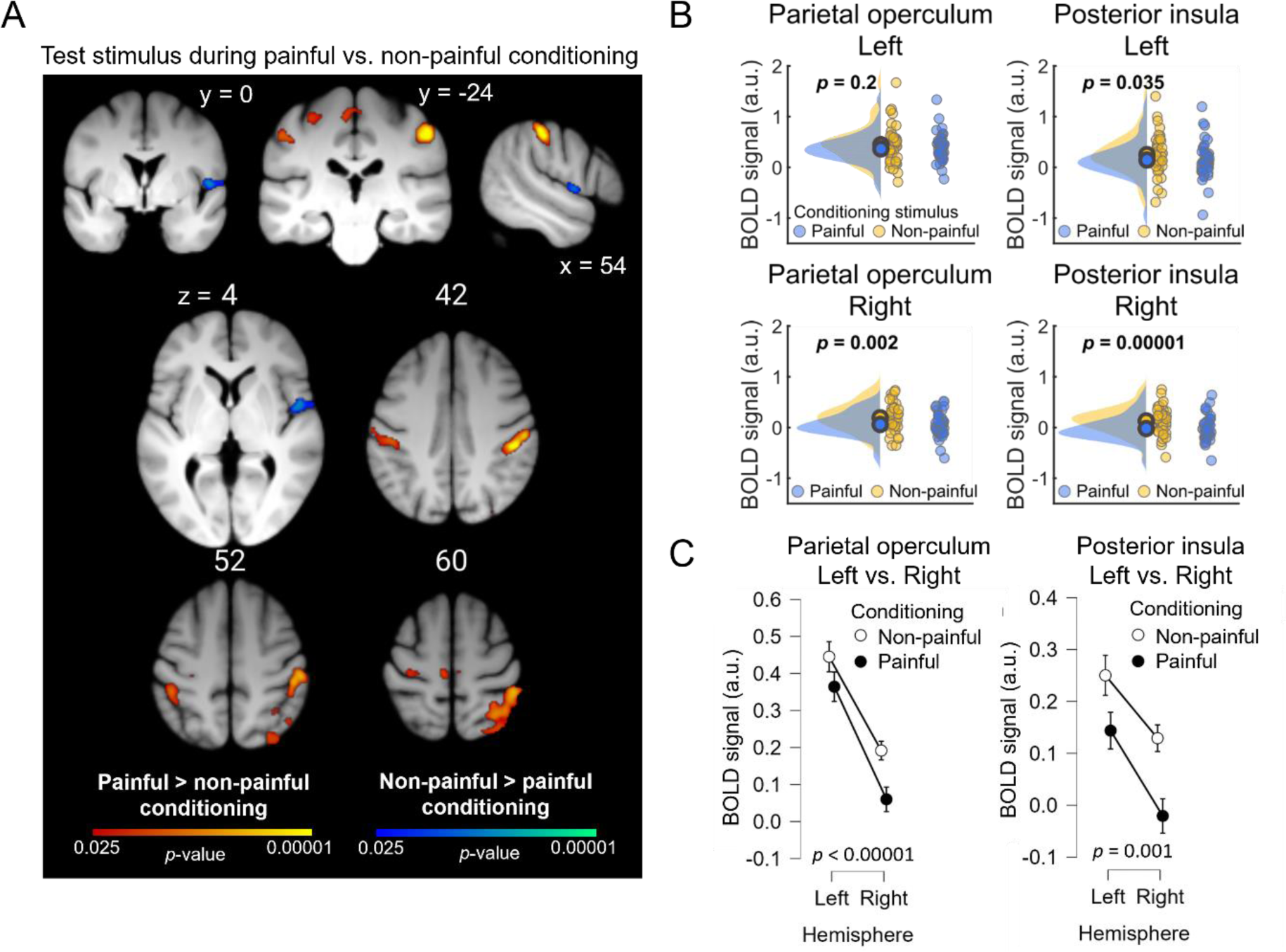
Brain BOLD fMRI activity associated with phasic test stimulus on the right arm in combination with either painful or non-painful conditioning stimulus (CPM). (**A**) Whole-brain analysis results with TFCE, FWE-corrected at *p* < 0.025 (corrected for two-tailed one-sample t-test). A region spanning between bilateral S1 and superior parietal lobule, left M1, and left anterior paracentral lobule showed increased activity when test stimuli were presented in combination with a painful in comparison with non-painful conditioning stimulus (“anti-CPM” effect). In contrast, decreased activity for test stimuli in combination with painful in comparison with non-painful conditioning stimulus was observed in right parietal operculum (“CPM” effect). (**B**) BOLD signal parameter estimates extracted from a priori defined anatomical regions-of-interest. Significant differences between parameter estimates for test stimuli presented during painful and non-painful conditioning stimuli were found in right parietal operculum and posterior insula (and at trend-level in left posterior insula), where activity was decreased for the test stimulus when it was presented in combination with painful rather than non-painful conditioning stimulus (one-tailed paired t-test, *p* < 0.0045 considered significant when correcting for 11 ROIs). (**C**) In both regions, the right hemisphere showed significantly lower BOLD signal compared to the left side while significant difference between conditioning stimulus intensities was only observed in the right hemisphere.

We also investigated if neural activity related to test stimuli was associated with the individual mean rated CPM magnitude or post-experiment verbal report (hypoalgesia, no difference, or hyperalgesia). We had an a priori hypothesis that higher subjective CPM (more hypoalgesia) would be associated with lower BOLD responses and we tested this in separate second-level covariate analyses, with TFCE at threshold *p* < 0.05 FWE-corrected (one-tailed paired t-test for). Higher mean rated CPM during the experiment was associated with lower BOLD responses to test stimuli overall, without interaction with conditioning intensity, in left anterior and posterior insula, left central operculum, left parietal operculum, middle cingulate cortex, left putamen, and left supramarginal gyrus; Supplementary Figure S2A). On the other hand, post-experiment verbally reported CPM was associated with lower BOLD responses to test stimuli over time during painful rather than non-painful stimuli in a small area in the right precuneus (Supplementary Figure S2B).

### Spinal cord activity related to phasic test pressure pain

We also hypothesized to observe increased BOLD activity correlated with test stimuli onsets on the right side of the spinal cord dorsal horn in segment C4, C5 or C6. We found significant BOLD response for test stimuli in right dorsal horn at segment C4 when averaging over all conditioning stimulus blocks, around coordinates x = 2, y = -46, z = -136 (‘Test stimulus during painful & non-painful conditioning > baseline’ contrast; TFCE FWE-corrected at *p* < 0.025; Figure 7A). We did not find any significant differences in BOLD activity to test stimuli during painful compared with non-painful conditioning. Comparison of mean BOLD signal estimates extracted from anatomical left dorsal horn ROIs for the three segments also did not show any significant differences in test stimulus responses between painful and non-painful conditioning stimuli, all *p* > 0.15. Because we had observed effects in the opposite-to-expected hemisphere or hemicord in our previous analyses, we also explored whether there would be any differences on the left side of dorsal horn but that was not the case, all *p* > 0.19. While no significant difference between conditioning stimulus conditions appeared across left and right dorsal horn at C4, *F*(1,41) = 2.00, *p* = 0.16, the BOLD signal was on average significantly higher on the right than on the left side of dorsal horn, *F*(1,41) = 24.2, *p* = 0.0001, 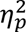 = 0.37, no interaction, *p* = 0.85. This effect seems to be driven by decreased BOLD responses on the left side (painful conditioning one sample t-test against zero, *t*(41) = −2.43, *p* = 0.02, *d* = 0.38, not corrected for multiple comparisons), rather than any clearly increased BOLD responses on the right side (painful or non-painful conditioning one-sample t-test against zero, both *p* > 0.4). All in all, we could not observe a CPM related signal difference in the spinal cord.

**Figure 7.**
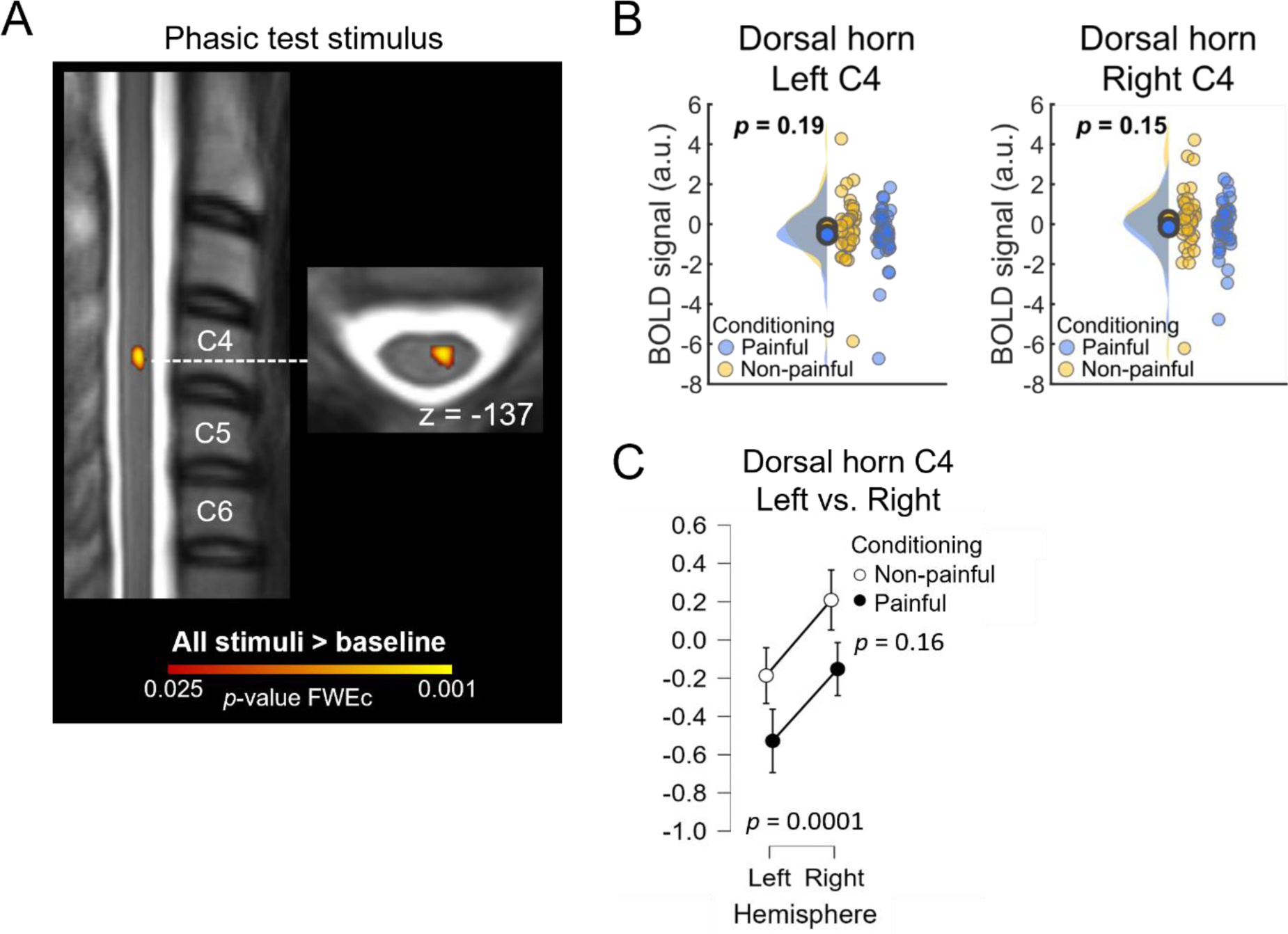
Spinal cord BOLD fMRI activity associated with phasic test stimulus on the right arm in combination with either painful or non-painful conditioning stimulus. (**A**) Spinal cord analysis with TFCE, FWE-corrected at *p* < 0.025 (corrected for two-tailed one-sample t-test) within right dorsal horn segments C4, C5 and C6. Test stimuli were associated with a significant activity in right dorsal horn at segment C4. A larger surrounding cluster presented at TFCE *p* < 0.001 uncorrected in blue for illustration. (**B**) BOLD signal parameter estimates extracted from a priori anatomical ROIs in the dorsal horn at segments C4, C5 and C6. There were no significant differences between mean BOLD signal estimates for test stimulus responses presented during painful and non-painful conditioning stimuli in right dorsal horn. We also did not find any differences on the left side in an exploratory analysis; therefore, there was no evidence for pain modulatory activity – CPM or “anti”-CPM – in the spinal cord. (**C**) Right side of the spinal cord dorsal horn had significantly higher BOLD signal to test stimuli than the left side dorsal horn but there was no significant interaction with conditioning stimulus intensity.

Similar to the brain, we also conducted analyses with rated CPM magnitude and verbal report of CPM as covariates. Unlike in the brain, there were no significant covariate effects for rated CPM magnitude in the spinal cord. However, post-experiment verbally reported CPM (hypoalgesia) was associated with higher test stimulus BOLD responses during both painful and non-painful conditioning in left dorsal horn at segment C5 (Supplementary Figure S2C).

### Brain-spinal cord connectivity changes for combined conditioning and test pressure

We had exploratory hypotheses of differences in functional connectivity (generalized psychophysiological interactions, gPPI) between nodes of the descending pain modulatory pathway from the brain (rACC) to brainstem (PAG) and medulla (RVM) to the spinal cord depending on conditioning stimulus (painful vs. non-painful). In a whole-brain analysis with TFCE at threshold *p* < 0.025 FWE-corrected (corrected for two-tailed paired t-test for both positive and negative connectivity changes), we observed decreased functional connectivity during conditioning stimuli in the cuneus, precuneus, and superior parietal and occipital regions (calcarine and lingual gyri) with a spinal seed region at segment C4 that was responsive to test stimuli on average (during painful or non-painful conditioning; Figure 7A). Importantly, this decrease in functional connectivity between the various parietal and occipital regions, including left hippocampus, and the spinal cord was not different between painful and non-painful conditioning (Figure 8A). Moreover, when investigating a gPPI in three of our central anatomical ROIs (rACC, PAG, and RVM), we found significantly increased functional connectivity during test stimuli between right spinal cord dorsal horn at segment C5 and rACC seed region (Figure 7B). Again, this effect was not different for painful and non-painful conditioning. Thus, we found evidence of decreased functional connectivity between sensory brain regions and the spinal cord in response to conditioning overall as well as increased functional connectivity between right C5 dorsal horn and rACC in the brain, but no correlate of CPM-like effects as the connectivity changes were not significantly different during painful and non-painful conditioning.

**Figure 8.**
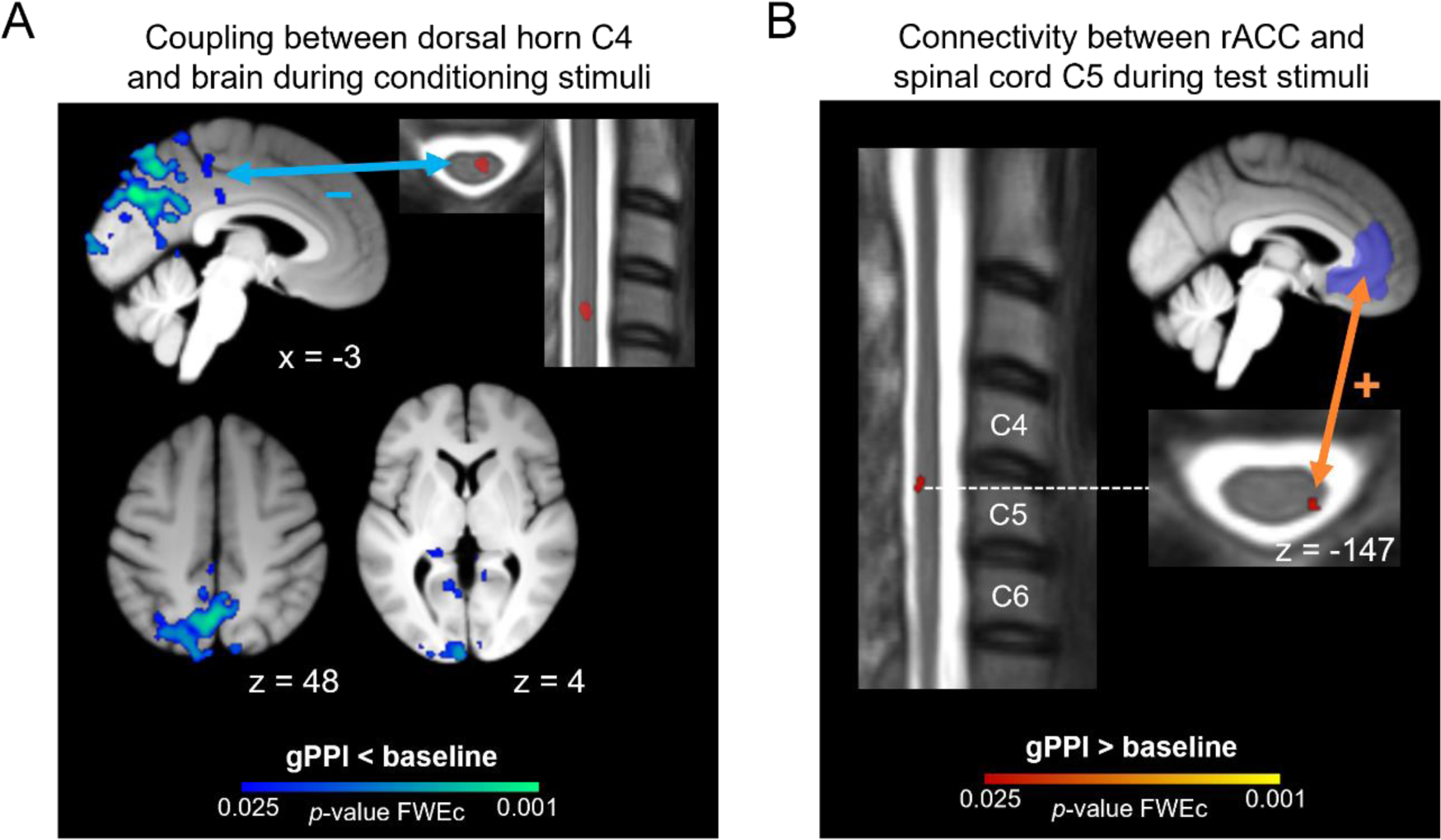
Neural coupling between brain and spinal cord, and brainstem and spinal cord. (**A**). Decreased functional connectivity (generalized psychophysiological interaction, gPPI) during conditioning stimuli in parietal and occipital regions and left hippocampus with seed region spinal cord dorsal horn C4 region responsive to test stimuli, regardless of conditioning intensity. (**B**) Increased gPPI functional connectivity during test stimuli between right dorsal horn at segment C5 and the anatomical seed region rACC in the brain, regardless of conditioning intensity. All results TFCE FWE-corrected at *p* < 0.025 (corrected for two-tailed one-sample t-test).

## Discussion

We investigated pressure pain responses and the modulation of these responses by concurrent conditioning pain in another limb across the cerebral cortex, brainstem, and cervical spinal cord in healthy humans. Subjective hypoalgesia during pain on both upper arms developed over repeated stimulation. Distinct neural responses to long-lasting conditioning pressure and short test pressure pain were found in pain processing brain regions as well as in the spinal cord dorsal horn in segments C6 and C4 respectively. Right parietal operculum and posterior insula exhibited decreased activity to short pressure pain on the left arm during painful conditioning compared with non-painful pressure. In contrast, bilateral primary somatosensory cortices and right superior parietal regions showed increased activity when the pressure stimuli were concurrently painful on both arms. Higher subjectively rated CPM was associated with decreased activity in pain and somatosensory regions including anterior and posterior insula and operculum. Moreover, we did not find BOLD signal changes related to conditioned pain modulation in the brainstem or the spinal cord. While neural coupling between the brain and spinal cord was altered during both long conditioning pressure and short pressure pain in comparison with baseline, the coupling strength was not associated with conditioning intensity.

### Development of hypoalgesic subjective and neural responses over time

The CPM effect we observed in trial-wise pain ratings developed over repeated application of test stimuli. Consequently, we did not observe an overall difference in rated CPM magnitude averaging over the entire experiment. While we observed very large inter-individual differences, individual mean CPM was not correlated with any background variables (age, sex, body mass index), individual parameters of pain calibration (e.g., calibrated test or conditioning stimulus intensity, or number of calibration stimuli), or ordering of conditions during the experiment (calibration arm order, first or last block conditioning intensity). Therefore, we are confident that our sample was well-balanced and the paradigm counter-balanced properly, and that the lack of a mean difference over all stimuli was indeed due to the necessity of factoring in the repeated stimuli.

In contrast to test stimuli, the painfulness of the conditioning stimulus was on average perceived as higher at the end of the experiment than in the beginning of the experiment. This supports CPM interpretation of our results since the ratings do not indicate the alternative possibility that the phasic but repeatedly applied painful pressure stimuli suppressed painfulness of the tonic pressure stimuli. We also did not find evidence for temporal summation of test pain, although it is possible that temporal summation existed physiologically but was cancelled out by CPM in the ratings. It may also be that temporal summation diminished the observed CPM effect but, in that case, less CPM would be expected at the end of the experiment when there would be most temporal summation – which was not the case. However, while conditioning stimuli were non-painful in two of the four experiment blocks, they were applied for 200 seconds at a time and the two conditioning stimulus trials were separated by only 30 seconds, leading to almost constant high pain for close to 7 minutes at a time. It is therefore possible that temporal summation underlies the observed increase in conditioning pain over time. Painfulness of the tonic conditioning stimulus may have also been increased over time by cumulative ischemic effects as high pressure applied as the cuff obstructs blood flow to the rest of the arm [30]. This caused short term “pins-and-needles” sensations and numbness in most participants as reported verbally after the experiment. Participants also reported that these sensations were subjectively aversive.

Taken together, participants experienced sensitization to the tonic pressure pain on the left arm while pain-inhibitory mechanisms were engaged for the phasic test pain on the right arm, and we do not think the effect can be explained by temporal summation of test pain but may be in part contributed to by summation of conditioning pain over time. On the other hand, pain modulation effects we observed in the brain were not significantly associated with the number of test stimuli. This difference between the behavioural and neural responses may reflect primacy of the modulation of the neural signals in relation to the behavioural painfulness ratings. It may also be that the statistical power to observe such an interaction effect in the BOLD signal was too low. Painful and non-painful conditioning blocks were counter-balanced across participants and therefore many participants experienced alternating blocks, which may have also reduced the cumulation of the CPM effect at least perceptually.

### Pain modulation in the cortex

Neural activity in the parietal operculum and posterior insula in response to test stimuli was decreased by painful conditioning pressure and may reflect inhibited signalling at the spinal level that propagates to the brain via ascending pathways, or alternatively top-down modulation of these sensory processing regions by cortical regions such as the ACC [4]. The BOLD signal in the parietal operculum and posterior insula was higher in the left hemisphere but the difference between conditioning intensities (non-painful > painful) was only significant in the right hemisphere. Since the test stimulus was applied on the right arm and the conditioning stimulus on the left arm, we expected neural responses to the test stimulus to appear preferentially in the left hemisphere, but the modulation was in the hemisphere contralateral to the conditioning stimulus. Taking this pattern of results together, it may be that left hemisphere was indeed more responsive to the test stimulus overall, but responses were still observed bilaterally and the concurrent stimuli on both arms led to a ceiling effect on pain modulation on the left but not the right side. Alternatively, the difference between painful and non-painful conditioning conditions that is seen in the right hemisphere might be a contralateral response to the conditioning stimuli since conditioning was on concurrently during all test stimuli. In the latter case, the effect may not precisely reflect CPM as the observed results would be due to the inability of our statistical model to distinguish test and conditioning stimulus related neural responses from each other because of their concurrent application - despite modelling the stimulus onsets separately.

We used parallel stimulation that has been posited to be a requirement for any CPM effect to reflect a DNIC-like mechanism in humans [33]. However, our protocol included homotopic stimulation where spinal mechanisms may be at play via communication within or across proximal segments, and only protocols with heterotopic stimulation can confirm a supraspinal CPM mechanism [28,33]. In principle, a neural CPM effect would be expected to be stronger contralateral to the test stimulus when looking at the brain and ipsilateral to the test stimulus when looking at the spinal cord. However, neural activity in response to pain is often bilateral due to cross-hemispheric communication [34]. Since we observed stimulus-ipsilateral pain modulatory responses in the brain, we cannot definitively conclude that these reflect neural correlates of ‘classical’ CPM although the behavioural evidence speaks for a CPM effect. On the other hand, we found that subjective rated CPM was associated with decreased BOLD signal in a wide range of somatosensory and pain processing regions in the left hemisphere at test stimulus onset, which supports the interpretation that neural correlates of CPM were present in the left hemisphere as well but did not reach significance in our main analysis.

### No pain modulatory responses in the spinal cord

Mirroring what was observed in the brain, much stronger neural signal at test stimulus onset was observed on the right side of the spinal cord dorsal horn at segment C4 than on the left side, as expected due to no crossing of sensory and motor fibres at the level of the spinal cord. The effect for conditioned pain modulation was in the expected direction (non-painful > painful conditioning) but it did not reach significance on either side of the cord. However, while the BOLD signal parameter estimates extracted from our anatomical ROI in dorsal horn of segment C4 were significantly higher on the right side compared to the left, they were in fact not significantly different from zero on the right side whereas the response during painful conditioning (but not non-painful conditioning) was significantly lower than zero, indicating decreased BOLD signal in comparison to baseline. Thus, we could not find evidence for BOLD signal difference in the spinal cord congruent with a CPM-like effect.

However, we would like to point out that our lack of evidence for CPM-induced decreased responding in the spinal cord does not mean that such an effect does not exist. There are a range of reasons why we might not have observed an effect. Firstly, it may be that our paradigm was not optimal, or the sample size was inadequately powered for spinal cord data compared to the brain. Visual observation of our BOLD signal estimates gives some indication that the signal variability was indeed larger in the spinal cord than in the brain (Figure 4, Figure 7). Secondly, due to the challenges inherent to spinal cord fMRI such as high impact of movement, poorer temporal signal-to-noise ratio, and higher susceptibility to distortions and signal loss [29], the signal quality in the spinal cord was worse than in the brain.

### Neural coupling during pressure (pain) across pain modulatory pathways

We found decreased neural coupling between brain regions including sensory occipital and parietal areas as well as left hippocampus, and the spinal cord dorsal horn C4 region responsive to test pain during conditioning compared with baseline, regardless of conditioning intensity (painful or non-painful). Since functional connectivity is not a directional measure, this may reflect either descending inhibition of the test stimulus responding in the spinal cord during the conditioning stimulus, or ascending competition of the two concurrent stimuli with higher test stimulus responding in the spinal cord leading to lower brain response to the conditioning stimulus. Moreover, we observed increased coupling between right dorsal horn at segment C5 and rostral ACC in the brain in response to test stimuli, again regardless of conditioning intensity. Since the rACC is an important region for pain modulation [4], the increased functional connectivity with the dorsal horn may reflect descending modulation of test stimulus responses in the spinal cord by rACC due to concurrent conditioning – but not exclusively by painful pressure. Unfortunately, we did not find any significant neural responses or connectivity changes in the brainstem, and so a node of the descending pathway required for signals from rACC to reach spinal cord is not accounted for in our data. Taken together, we observed functional interactions between neural activity in the brain and spinal cord, but the evidence does not support any CPM related connectivity changes during painful and non-painful conditioning.

### Limitations

Attention is known to modulate pain [35,36]. While attention to the conditioning stimulus can increase CPM and also produce hypoalgesia in addition or independent of CPM [37–40], a limitation of our study is that we cannot separate conditioning stimulus effects due to painfulness and attention or salience since we applied the conditioning and test stimuli in parallel [33,41]. However, we included a non-painful control condition to somewhat counter this and to be able to claim that our effects are not due to pressure per se but due to painful pressure. Regardless, the highly painful conditioning stimulus was more salient than the non-painful stimulus.

While participants were not informed about the number or ordering of the painful and non-painful conditioning blocks, since the experiment consisted of two painful and two non-painful blocks, participants could reasonably expect what was going to happen either in the second last or last block depending on the blocks they received before. Therefore, they may have been able to prepare for pain or relax in anticipation of relief from pain, and it is known that expectations can strongly influence perceived pain [35,42].

Unlike often recommended in discussions of CPM methodology (Piché, 2009; Yarnitsky, 2015, Sirucek et al., 2022), we did not use heterotopic stimulation as it was not possible to acquire fMRI signals from the spinal segment corresponding to the leg in combination with the cervical spinal cord and the brain. The choice of both upper arms allowed us to image the entire volume of interest within a repetition time of 1.9 seconds as the corresponding spinal segments are located closer to the brain. Naturally, this choice means that we could not have definitively excluded spinal modulation of noxious inputs in our spinal cord results but this is a moot point as we did not observe CPM-like effects in the spinal cord. Moreover, the spinal cord responses to conditioning pain and test stimuli were separated by at least one segment of the spinal cord.

We chose to use parallel test and conditioning stimuli because it is not precisely known how long CPM effects last [43] and it is likely dependent on various factors such as test and conditioning stimulus intensity, duration, and modality [43,44]. In a sequential protocol, CPM effect induced by a single conditioning stimulus before test stimulus application might not last long enough to measure test stimulus neural responses reliably. This would also exclude measuring neural responses to the conditioning stimulus, which was of interest to us. The two designs have not been systemically compared but one study with cold pressor conditioning did not find a significant difference in CPM magnitude between a parallel and sequential protocol [45]. It is however possible that our paradigm included CPM carry-over effects when a painful conditioning block was followed by non-painful conditioning block since we kept the inter-block interval short due to time constraints.

### Conclusions

We found distinct neural representation of parallel test and conditioning pressure in the brain and the spinal cord, as well as modulation of behavioural and brain responses to test pain by concurrent conditioning pain. The neural effects of “pain inhibits pain” (a CPM-like effect) and pain summation were represented in different brain regions including most somatosensory and pain processing regions and decreased neural activity in some of these regions was associated with higher subjective CPM. On the other hand, we could not find significant fMRI signals related to CPM in the brainstem or the spinal cord. Neural coupling between the brain and spinal cord was altered during both conditioning and test pain regardless of conditioning intensity. Our study included a relatively large sample size and a carefully balanced design, trial-wise pain ratings and control conditioning stimulus that have not been included in previous studies, as well as imaging across almost all relevant parts of the descending pain modulatory pathway. Future directions to pinpoint the neural mechanisms of CPM could include studies with heterotopic stimulation that focus investigations on test stimulus responses with and without painful conditioning in the cervical spinal cord to confirm supraspinal modulation of spinal cord activity. Developments in corticospinal imaging, with sequences better optimized for brainstem and spinal cord image quality and signal-to-noise ratio, may be required to detect CPM effects across the different levels of the pain modulatory pathway.

## Materials and Methods

### Pre-registration

The study was pre-registered at the German Clinical Trials Register, and the pre-registration can be accessed at: https://drks.de/search/en/trial/DRKS00027882.

### Participants

Participants were recruited through advertisements on an online platform, leaflets, and word-of-mouth and screened for eligibility based on the following criteria: generally healthy, right-handed, age between 18 and 45 years, body-mass-index (BMI) between 18 and 30, no contraindications to MRI, no acute or chronic pain, no acute or chronic somatic or psychiatric illness, no recent or chronic injury in the arms, no current muscle soreness or large bruising in the arms, and no regular intake of medication (except for contraceptive pill, allergy medication, or medication for hyper- or hypothyroidism). Participants were asked to refrain from intensive sports on the day of the experiment and to not take pain killers in the 24 hours preceding the experiment. The study was approved by the Ethics Committee of the Medical Council of Hamburg (2020-10144-BO-ff) and conducted according to the Declaration of Helsinki. Written informed consent was be obtained from all participants prior to study participation, and after the study, participants were compensated with 45-55€ depending on study duration (usually 2 hours).

In total, 50 participants took part in the study, out of whom 43 completed the study according to protocol (4 participants stopped the experiment due to anxiety or too high pain sensitivity; 1 participant fell asleep during the experiment; 2 participants were excluded due to technical issues). After excluding 1 participant who had extremely low rating of the conditioning pressure pain that was intended to be painful, the final sample was 42 participants (21 women, 21 men, mean age 26.7 ± 5.3 years, mean BMI 23.7 ± 3.2 kg/m^2^).

### Experimental design

After arrival at the laboratory, participants filled out consent forms, MRI safety forms, and psychological questionnaires on attitudes toward pain. Next, participants underwent a conditioned pain modulation experiment consisting of calibration, pre-experiment conditioning stimulus painfulness rating (with Visual Analog Scale, VAS), four blocks of conditioned pain modulation experiment, and post-experiment conditioning stimulus painfulness rating inside the MRI scanner (Fig. 1A). Finally, participants answered verbal questions relating to the experiment and were debriefed.

#### Stimulus presentation

The conditioning stimulus was a tonic pressure stimulus on the left arm and the test stimulus was a phasic pressure stimulus on the right arm (Fig. 1B), delivered with computerized cuff algometry (CPAR device, Nocitech, Denmark). The conditioning stimulus was continuous pressure for 200 s at a level that was either non-painful (set to fluctuate between 2 and 5 kPa pressure output for each participant) or moderately-to-highly painful (individually calibrated to fluctuate between 50 VAS trough and 70 VAS peak intensity). The non-painful control and painful experimental conditioning stimuli were presented in separate blocks (two control blocks, two experimental blocks). Each block consisted of two conditioning stimuli of 200 s duration with a sinusoidal cyclic form where peaks and troughs were repeated three times. The test stimuli lasted 5 s and were always presented at the same moderately painful level (individually calibrated to around VAS 60). Since the test pressure duration was always 5 s, those participants with higher calibrated pressure levels (kPa) for the specified painfulness level had a faster initial rise-time of the stimulus. The experiment was programmed with Psychtoolbox 3.0.17 (psychtoolbox.org) [46] and MATLAB2020a (The MathWorks) and presented from a Windows computer with MATLAB2016 to the participants from a screen behind the scanner with the image reflected to the participants eyes via a mirror inside the scanner.

#### Pressure stimulus calibration

Calibration of the pressure stimuli took place with the participant lying inside the MRI scanner while preparatory scans were obtained to save time and to have a similar environment for calibration as for the experiment. Tonic conditioning stimuli (left arm) and phasic test stimuli (right arm) were calibrated separately. Out of the final sample, 22 participants underwent calibration procedure for conditioning stimuli first and test stimuli second, and 20 participants did the procedure the other way around. First, participants received two low pressure stimuli (10 and 20 kPa) to get used to the feeling of the pressure cuffs inflating. Next, a pain threshold search was conducted where participants started with pressure at usual pain threshold based on pilot studies and rated in total six stimuli as either non-painful or painful. Lowest pressure rated as painful was taken as the pain threshold. The pain threshold search was completed for both arms before participants were familiarized with the VAS painfulness scale and continued on to be exposed to and rate on the VAS four stimuli above the pain threshold with increasing pressure, except for the fourth one which is the same as the second one in order to check if the participant’s perception of the intensity changed depending on how high intensities they were already exposed to. Based on this rating data, a linear regression algorithm calculated target VAS 10, 30 and 90 pressures that the participant rated on the VAS, and these data points were added to the regression to get improved estimates for the final step of calibration. In the final step, less covered bins of the painfulness scale from 0 to 100 were filled in with new ratings from the participant and the regression was adjusted. This continued until enough coverage was reached to reliably estimate the required pressure for the pain level (in VAS units) for the experiment (VAS 60 for test stimuli/right arm, VAS 50-70 for conditioning stimuli/left arm). The entire procedure after familiarization with the VAS was completed for each arm/stimulus type in a counter-balanced manner across participants.

#### Experiment

Participants saw a grey background with a white cross in the middle for fixation except during rating periods. Firstly, participant experienced 80 seconds of the conditioning stimulus (one cycle) to rate the intensity of the stimulus before the experiment. It was found during pilot testing that participants most often reached a plateau in their conditioning stimulus rating after around 60 seconds and we did not want to expose participants to too long painful stimuli before the experiment, especially considering the extensive calibration for two arms. Next, four blocks of the conditioned pain modulation experiment followed with the order of the control and experimental blocks counterbalanced across the participant sample. During the experiment, the conditioning stimuli were on for 200 seconds, during which nine test stimuli would come on for 5 seconds each, with a 5-second painfulness rating period taking place immediately after the offset of the test stimulus. The onsets of the test stimuli were jittered (±0-2 s) around defined timepoints during each cycle of the conditioning stimulus (at 5 s or 20 s post-onset on the upward slope toward the peak, or at 35 or 50 s post-onset on the downward slope toward the trough), with the nine test stimuli randomized to three points on each of the three cycles of the conditioning stimulus. The painfulness ratings were done on a scale of 0-100 where 0 was instructed to be non-painful, and 100 to be pain that would be intolerable for an extended duration in the context of the experiment. The inter-stimulus interval between successive test stimuli was 8-37 seconds. The inter-trial interval between the two conditioning stimuli of each block was 30 seconds and the inter-block interval was 60 seconds. In total, there were four control conditioning stimuli and four experimental conditioning stimuli, and a total of 72 test stimuli split between the two conditioning intensity conditions (36 + 36). In the final part, participants repeated the 80-seconds rating of the conditioning stimulus.

#### Physiological measures

Respiration and pulse were measured throughout the entire experiment. Pulse was recorded with photoplethysmogram clip from the big toe of the left foot and respiration with and a breathing belt, both via Expression system (In Vivo, Gainesville, USA). Physiological signals, scanner pulses as well as stimulus timing markers sent from the MATLAB experiment program were monitored and digitized with a CED1401 system and saved with Spike 2 software (Cambridge Electronic Design) on a Windows computer. The physiological log files were imported in MATLAB, and respiration and pulse signals were downsampled to 100 Hz, detrended and smoothed (pulse with 5 and respiration with 50 samples FWHM Gaussian kernel).

#### Post-experiment questions

After the experiment, participants were asked the following questions verbally by the experimenter: A. “When you compare the parts of the study where you received: 1) moderate-to-high painful long pressure and 2) low, non-painful long pressure on the left arm, did you feel any difference in the painfulness of the short painful stimuli on the right arm? (yes/no)”, and B. “If yes, the short stimuli during high long pressure were… (more painful/less painful) …than during low long pressure”. Participants were also asked about any side effects or other sensations they may have experienced during the experiment relating to the pressure on the arms.

### MRI data acquisition

All MRI acquisitions were performed on a 3 Tesla whole-body MR system (Magnetom PrismaFit, Siemens Healthineers, Erlangen, Germany) using a standard 64-channel head-neck coil. Participants were be positioned with their chin at the level of the isocentre. During the experiment, T_2_*-weighted echo-planar imaging (EPI, flip angle 70°, TR 1991 ms, descending slice acquisition order) covering 60 slices without gap in the brain (FOV 220 x 220 mm^2^, voxel size 2.0 x 2.0 x 2.0 mm^3^, TE 24 ms, 56.0 ms per shot) and 12 slices without gap in the spinal cord (FOV 132 x 132 mm^2^, voxel size 1.2 x 1.5 x 5.0 mm^3^, TE 27 ms, 72.6 ms per shot). Simultaneous multi-slice imaging (blipped-CAIPI) was applied for the brain slices with an acceleration factor of 3 [47]; slice-specific z-shim [48], and a saturation pulse covering the throat and chin region were applied for the spinal cord slices. Adjustment volumes covered the full brain EPI volume and a 40 x 40 x 60 mm^3^ volume focusing to the spinal cord, respectively. The brain field of view was aligned with the bottom-most slice including rostroventral medulla below the pons and the cerebellum in its entirety. Consequently, in some participants the top of the brain including primary somatosensory cortex was not fully included. For the spinal cord, the topmost slice was positioned to the upper edge of the spinal disc C3-C4.

Anatomical images included a high-resolution structural T1-weighted scan covering the entire head and the neck up to the lower part of the second or upper part of the third thoracic vertebra, depending on participant height (MPRAGE; TR 2300 ms, TE 3.41 ms, inversion time 1100 ms, 1.0 x 1.0 x 1.0 mm^3^ voxel size, flip angle 9°, FOV 320 x 240 x 192 mm^3^, sagittal slice orientation), and a T2-weighted acquisition of the spinal cord including vertebrae C2-T2 (3D turbo spin echo, TR 1500 ms, TE 120 ms, 0.8 x 0.8 x 0.8 mm^3^ voxel size, refocusing flip angle 120°, FOV 256 x 256 x 51 mm, 64 sagittal slices).

After a localizer acquisition, EPI volumes and the corresponding adjustment volumes were positioned, and shim settings for the fMRI experiments were determined using a standard field map acquisition and a dedicated shim algorithm as described recently [49]. The time to calculate the shim settings (static and dynamic updates for the brain volume) was used to acquire the T1-weighted anatomical images. It was followed by the z-shim reference acquisition covering the spinal cord volume only, during its analysis the T2-weighted acquisition was performed. Finally, an EPI acquisition with z-shim was performed to check the determined values before the fMRI experiments were started.

### MRI data preprocessing

#### Brain preprocessing

After acquisition, DICOM files were converted into the NIfTI files, separately for the brain and the spinal cord sub-volumes. Preprocessing and statistical analyses were also conducted entirely separately for the brain and spinal cord subvolumes. Preprocessing of the brain fMRI data was done fully in SPM12 (Wellcome Trust Centre for Neuroimaging, London, UK) and included slice timing correction, realignment, and non-linear co-registration. Since we wanted to correct the slices for timing difference between brain and spinal cord subvolume acquisitions, onset slice of slice-timing correction for brain was set to the last brain slice that was closest to the spinal cord. First-level general linear models (GLMs) for each participant included individual brain masking (white matter and grey matter mask from segmented T1 smoothed with 6 mm FHWM kernel, excluding skull and eyes and some parts of cerebrospinal fluid), bandpass filtering to remove low-frequency drifts with cycles longer than 200 s, and autocorrelation modelling with ‘FAST’. Motion correction was done by including 24 motion parameters (derivatives of the initial 6 motion parameters from realignment, and squares of the original and derivative parameters) in the first-level GLM as nuisance regressors. Next to the motion parameters, the first-level nuisance regressors also included 18 RETROICOR regressors (4 orders of cardiac, 3 orders of respiratory and 1 order of interaction components) [50,51], and 6 principal components each for noise regions including white matter (WM), cerebrospinal fluid (CSF), and the interface of white matter and cerebrospinal fluid [52]. The physiological noise regressors were calculated with the TAPAS PhysIO toolbox version 8.1.0 [53]. Settings for creating the noise region masks were as follows: WM and CSF threshold 0.7, erosion level 1; CSF x WM threshold 0.15, erosion level 0. Motion censoring was performed by excluding volumes with more than 2 mm translation or 2° rotation and the motion censored volumes were included in the first-level GLM as nuisance regressors (1 regressor per censored volume). Spatial normalization to a the MNI template and smoothing with 6 mm kernel was done on contrast images from the first-level GLMs, and these normalized and smoothed contrast images for each participant were taken to the second-level analysis.

#### Spinal cord preprocessing

Spinal cord data was preprocessed with the Spinal Cord Toolbox version 5.6 [54]. Since we wanted to approximately correct the slices for timing difference between brain and spinal cord subvolume acquisitions, onset slice of slice-timing correction for spinal cord was set to the first spinal slice that was closest to the brain. The anatomical T2 image of the spinal cord for each participant was segmented with a deep learning model (the 3D option was used for most participants but 2D was used for a couple of participants when 3D failed) [55]. Next, the vertebrae were labelled automatically with the help of the segmentation, and the T2 images registered to the PAM50 template space [56] using the segmentation as well as the vertebrae labels (default settings). The resulting 4D warping field was saved for later purposes. Next, mean functional image for each scanner run was calculated and spinal cord segmentation applied to these mean EPIs as for the T2 images. T2 images were coarsely registered to the mean EPIs (1-step ‘sct_register_multimodal’ with parameters type ‘seg’ [segmentation], algorithm ‘affine’, metric ‘MeanSquares’, and smooth ‘10’ mm), and these co-registered T2 images were segmented in order to create a mask of the spinal cord in EPI space. Individual EPIs were then corrected for motion with SCT’s algorithm, using the spinal cord mask to limit the voxels considered by the algorithm. Notably, SCT’s motion-correction algorithm includes only six degrees of freedom and does not consider translation in z-direction (slice-wise movement). However, we found the motion correction results satisfactory and additionally included the 24 brain motion parameters, as well as 18 RETROICOR regressors, in the later first-level GLM design as nuisance regressors (assuming that z-direction motion of the spinal cord slices would be highly correlated with the z-direction motion of the brain slices). Moreover, we included censoring regressors for those volumes with excessive motion (> 2 mm translation or 2° rotation; plus a few additional volumes detected by eye). Motion-corrected mean EPIs were then segmented for a better final segmentation result, and precisely co-registered to the T2 images (3-step ‘sct_register_multimodal’ where parameters were: step 1) type ‘seg’, algorithm ‘translation’, metric ‘MeanSquares’, and smooth 20 mm; step 2) type ‘seg’, algorithm ‘affine’, metric ‘MeanSquares’; step 3) type ‘im’ [image], algorithm ‘syn’ [Symmetric image normalization], metric ‘MI’ [Mutual Information], iterations ‘5’). A combined warp field including the inverse of EPI warp to T2, and T2 warp to PAM50 template were used to normalize the EPI-co-registered T2 images to the PAM50 template (resolution 0.5 x 0.5 x 0.5 mm). Normalization accuracy of the contrast images was visually inspected and deemed adequate around the regions of interest in the cord. Normalized images were cropped to exclude image space that was not part of the actual imaged volume. First-level general linear models (GLMs) for each participant included masking with PAM50 template binary mask of the spinal cord, bandpass filtering to remove low-frequency drifts with cycles longer than 200 s, and autocorrelation modelling with ‘FAST’. Finally, contrast images were smoothed with [2 2 3] mm FWHM Gaussian kernel before entering them into the second-level group analysis.

### fMRI analyses

#### Whole-brain and whole-cord statistical analysis

First-level GLM design matrices for both brain and spinal cord subvolumes were built with fMRI runs/experimental blocks concatenated as one session (4 x 227 = 908 volumes). The design matrices included experimental condition regressors ‘test stimulus onset’ (modelled as a 5-s boxcar function) and ‘conditioning stimulus onset’ (modelled as two zero duration stick functions per TR, i.e. ∼1 stick per 1 s, for a total duration of 200 s). Separate onset regressors were included for painful and non-painful conditioning stimulus blocks except for one ‘pain rating onset’ regressor, totalling 5 onset regressors. Session constants were added to the end of the design matrix separately for each run. Additionally, both ‘test stimulus onset’ regressors had a parametric modulator regressor ‘stimulus index’, which was defined as numbers 1-72 representing each test stimulus, mean-centered and split for the two conditioning stimulus conditions and added after their respective onset regressor in the design matrix, to test for time effects across the entire experiment. We also included parametric modulators ‘conditioning pressure’ (200 s cyclical shape without intensity information across blocks) and ‘conditioning pressure x test pressure’ for the ‘conditioning stimulus onset’ regressor to investigate if there was any effect of the conditioning stimulus sinusoidal shape and/or interaction effect of the conditioning and test stimulus shapes, but these analyses are not reported further as they were very exploratory and did not show any promising results. Nuisance regressors and temporal filtering were included as described above in the preprocessing sections and experimental regressors were convolved with the canonical haemodynamic response function (HRF) in SPM. All experimental and nuisance regressors were z-scored and SPM explicit orthogonalization for parametric modulators was turned off. After model estimation, the following first-level contrasts were defined for each participant: 1) ‘conditioning stimulus onset non-painful > baseline’, 2) ‘conditioning stimulus onset painful > baseline’, 3) ‘conditioning stimulus non-painful & painful > baseline’, 4) ‘conditioning stimulus painful > non-painful’, 5) ‘test stimulus onset during non-painful conditioning stimulus > baseline’, 6) ‘test stimulus onset during painful conditioning stimulus > baseline’, 7) ‘test stimulus during painful & non-painful conditioning stimulus > baseline’, 8) ‘test stimulus during painful > non-painful conditioning stimulus’, 9) ‘test stimulus onset x stimulus index during non-painful conditioning stimulus > baseline’, 10) ‘test stimulus onset x stimulus index during painful conditioning stimulus > baseline’, 11) ‘test stimulus onset x stimulus index during non-painful & painful conditioning stimulus > baseline’, 12) ‘test stimulus onset x stimulus index during painful > non-painful conditioning stimulus’, and 13) ‘pain rating onset > baseline’.

The contrast images from the first-level analysis were entered into a second-level GLM separately for brain and spinal cord subvolumes creating one one-sample t-test model for each of the 13 first-level contrasts. Second-level brain mask was set as the thresholded at 0.8, 2 mm FWHM smoothed, skull-stripped mean anatomical image of all participants, and second-level spinal mask was set as normalized PAM50 spinal cord template resliced into the EPI space. Second-level models were estimated with traditional SPM approach, but for statistical thresholding, we used Threshold-Free Cluster Enhancement (TFCE) [57] as implemented in Christian Gaser’s TFCE toolbox for SPM (https://neuro-jena.github.io/software.html) with the following default settings: 5000 permutations, step size 100, nuisance method 2 (Smith if nuisance regressors found, otherwise Draper-Stoneman), extent exponent 0.5 (more weighting of local effects), and height exponent 2 as recommended in [57]. For all analyses, statistical inferences were made based on “TFCE_log_pFWE” files with threshold p < 0.05, that is, ±log10(0.05) = ±1.301 for one-tailed comparisons and ±log10(0.025) = ±1.6021 for two-tailed comparisons.

#### Region-of-interest (ROI) analysis

Our anatomical a priori ROI masks for the brain volume included bilateral rACC, bilateral vmPFC, bilateral dlPFC, right and left anterior insula, right and left posterior insula, right and left parietal operculum (SII), bilateral PAG, and bilateral RVM. All of these ROIs, except parietal operculum (due to accidental omission) were included in the ethics proposal and pre-registration of the study and masks finalized before looking at the main effects of interest (CPM-related effect). Rostral ACC mask was created by combining Area s24, Area p24ab, Area p24c, and Area s32, vmPFC by combining Fo1 and Fo2, and dlPFC by combining MFG1 and MFG2 from the Jülich Probabilistic Atlas [58] with SPM Anatomy Toolbox version 3.0 [59], including both hemispheres. The dlPFC mask was additionally extended with a 6 mm radius sphere centred around coordinate 48, ±28, 24 from a previous CPM fMRI study [60] to include a more posterior part of the dlPFC that was not included in any of the Jülich atlas regions. Anterior insula mask was created by combining Id4, Id6, and Id7, (dorsal and inferior) posterior insula mask by combining Ig1, Ig2, Ig3, Id1, Id2, Id3, Id5, and Ia1, and parietal operculum by combing OP1, OP3, and OP4 of the Jülich Probabilistic Atlas, separately for the two hemispheres. The probabilistic atlas masks were binarized with threshold > 0, smoothed by [3 3 3] mm FWHM Gaussian kernel and binarized once more. Finally, overlap between the three prefrontal masks was removed by excluding vmPFC mask voxels from rACC mask, and rACC mask voxels from dlPFC mask, and each mask was also masked with the second-level brain mask (smoothed mean anatomical image of all participants) to ensure that all voxels of the masks were within the volume of interest. PAG and RVM masks were obtained from the Open Science Framework repository for [61] (https://osf.io/xqvb6). Since we had the a priori expectation that the relevant activity in the spinal cord should be limited to the dorsal horn in segments C4, C5 or C6, these were chosen as our anatomical ROIs for the spinal cord volume. The anatomical masks were defined as spinal level 5 (corresponding to spinal segment C4), spinal level 6 (C5), and spinal level 7 (C6) from the PAM50 template.

Individual beta parameter estimates for each relevant condition (test stimulus and conditioning stimulus onsets separately for non-painful and painful conditioning stimulus blocks) were extracted from each ROI with MarsBaR toolbox for SPM, version 0.45 [62]. Our comparisons of interest for the ROIs were: 1) ‘painful conditioning stimulus > non-painful conditioning stimulus’ (one-tailed t-test), 2) ‘teststimulus during painful vs. non-painful conditioning stimulus’ (two-tailed test), and 3) ‘test stimulus x stimulus index during painful vs. non-painful conditioning stimulus’ (two-tailed test). All p-values were Bonferroni-corrected for the number of ROIs, which was 11 for the brain volume (as we did not have hemisphere-specific hypotheses and wanted to test separately for both left and right sides for anterior insula, posterior insula, and parietal operculum) and 3 for the spinal cord volume (due to stimulus-ipsilateral left/right hypotheses). However, to formally test for differences between stimulus-ipsi/contralateral sides in the brain and spinal cord, we also conducted a 2×2 repeated-measures ANOVA with factors ‘hemisphere/cord’ (left/right) and ‘conditioning stimulus condition’ (painful/non-painful) for the following ROIs: anterior insula, posterior insula, parietal operculum, dorsal horn level 5, dorsal horn level 6, and dorsal horn level 7.

#### Covariate analyses

We included two second-level whole-brain/cord covariate analyses in SPM separately for 1) rated CPM magnitude, computed as the mean difference between trial-wise pain ratings during painful and non-painful conditioning (more positive values indicating more hypoalgesia and more negative values indicating more hyperalgesia), and 2) post-experiment verbal report of CPM (1 = hypoalgesia, 0 = no perceived difference in test stimulus pain between painful and non-painful conditioning, -1 = hyperalgesia). We included the same contrasts as for the main whole-brain/cord analyses and conducted TFCE on the covariate contrast. We had the a priori pre-registered hypothesis that higher CPM should be associated with decreases in BOLD signal (negative correlation) and therefore we only corrected for FWE at *p* < 0.05 for the negative contrast direction.

#### Functional connectivity

Functional connectivity analysis with generalized psychophysiological interactions (gPPIs) [63], was performed to investigate the co-occurring BOLD activation in the brain and spinal cord. Brain PPI seed ROIs were selected to be PAG, RVM and ACC (defined as in the previously described ROI analysis) due to their central involvement in the descending pain modulatory pathway [2,4]. For spinal cord seed ROIs, we defined spheres of 2 mm radius around significant peak coordinates from the small volume corrected analysis of test stimuli and conditioning stimuli (‘test stimulus during painful & non-painful conditioning > baseline’ peak coordinate at x = 2, y = -46, z = -136; ‘painful conditioning stimulus’ > baseline at 0, -48, -174; ‘non-painful conditioning stimulus’ > baseline at -4, -45, -152; ’painful conditioning stimulus > non-painful conditioning stimulus’ at -1, -48, -174). Time series data was extracted from each seed ROI, with the time course data for the brain analyses extracted from the spinal seeds, and vice versa. The seed time series, stimulus onsets (separately for test and conditioning stimuli for painful and non-painful blocks), and interactions of the time course with the stimulus onsets were entered as regressors in a first-level analysis for each participant and each seed ROI. The first-level PPI analyses included the same nuisance regressors as the previously described brain and spinal cord analyses. For second level statistical analysis, we used TFCE with same parameters as for the previous whole-brain and whole-cord analyses.

## Code availability

Code for all analyses is openly available on GitHub: github.com/karitaojala/CPM-Pressure.

## Data availability

The project is published at Open Science Network with DOI 10.17605/OSF.IO/C9UTP. Anonymized first-level contrast images and group-level statistical parametric maps (SPMs) for all contrasts of interest as well as ROI masks and extracted parameter estimates are available at https://osf.io/46nuw. Other data including pain ratings, post-experiment verbal report of CPM, and background variable data are available at https://osf.io/pjxvc.

## Acknowledgements

We thank Kristian Hennings of Nocitech for his extensive help in setting up our experimental stimulation paradigm using the CPAR device. We also thank Björn Horing for discussions, help with various technical aspects and for sharing code, and Waldemar Schwarz for his central role as the MR radiographer during our data acquisition.

## Notes

**Competing interests** None.

https://doi.org/10.17605/OSF.IO/C9UTP

### Competing Interest Statement

The authors have declared no competing interest.

