## Supplementary information for "Conditioned pain modulation of pressure pain is associated with reduced activation in the parietal operculum and posterior insula"

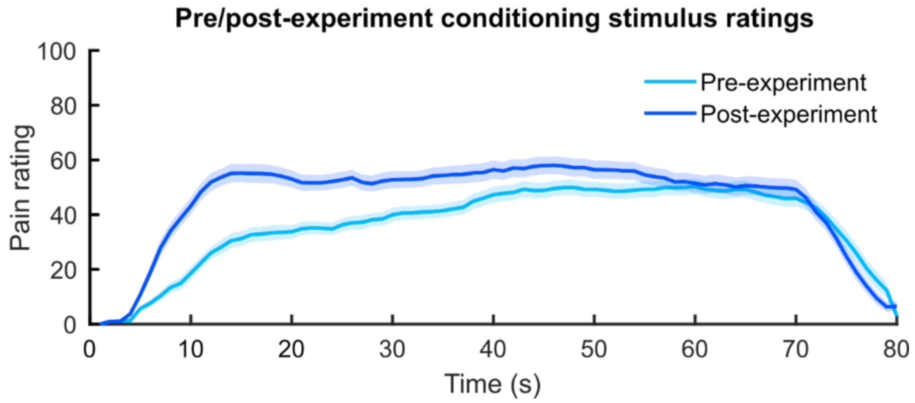


**Supplementary Figure S1. Pre- and post-experiment continuous pain ratings for conditioning stimuli.** Continuous pain rating for the tonic conditioning stimulus obtained right before and after the experimental blocks. The conditioning stimulus was rated for 80 seconds rather than the full 200 seconds to save time and not to induce too strong pain modulation already before the start of the experiment. Sensitization to the conditioning stimulus over time was observed, with higher pain ratings in the first ~55 s of the stimulus in the post-experiment vs. pre-experiment rating session.

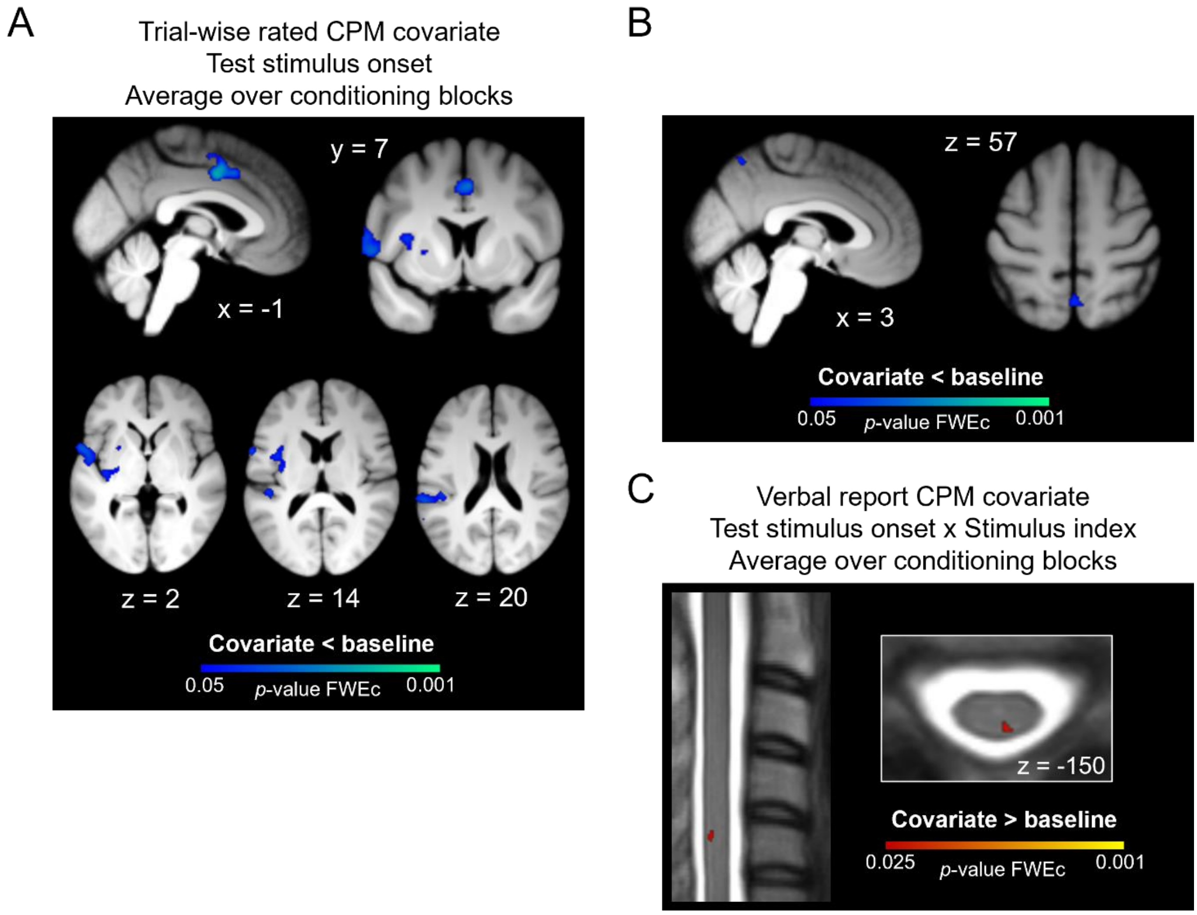


**Supplementary Figure S2. Whole-brain analysis with behavioural covariates mean rated CPM magnitude and post-experiment verbally reported CPM.**

(**A**) The mean difference between trial-wise test stimulus pain ratings during painful and non-painful conditioning was negatively associated with BOLD signals in left anterior and posterior insula, left central and parietal operculum, left putamen, left supramarginal gyrus, and middle cingulate gyrus.

That is, higher hypoalgesia of test pain during painful conditioning in comparison with non-painful conditioning was associated with decreased neural activity.
(**B**) Post-experiment verbal report of CPM was negatively associated with BOLD signal in the right precuneus, that is, reported hypoalgesia was associated with decreased neural activity while reported hyperalgesia was associated with increased neural activity.

**Supplementary Table S1. Correlation of average rated and verbal CPM with participants' background variables and experimental design's cross-balanced orders.**

| | | Spearman's rho | p-value |
|---|---|---|---|
| Rated CPM | - Sex (1 = F, 0 = M) | 0.051 | 0.748 |
| | - Age | -0.046 | 0.774 |
| | - Body mass index | -0.134 | 0.399 |
| | - First calibration arm (1 = left, 0 = right) | -0.026 | 0.872 |
| | - First block conditioning (1 = pain, 0 = no pain) | -0.267 | 0.087 |
| | - Last block conditioning | -0.004 | 0.980 |
| | - Test stimulus pressure | -0.257 | 0.101 |
| | - # calibration stimuli test pressure | 0.008 | 0.962 |
| | - Test stimulus calibration slope | 0.257 | 0.115 |
| | - Conditioning stimulus peak pressure | -0.234 | 0.136 |
| | - Conditioning stimulus trough pressure | -0.242 | 0.123 |
| | - # calibration stimuli conditioning stimulus | -0.219 | 0.181 |
| | - Conditioning stimulus calibration slope | 0.045 | 0.786 |
| Verbal CPM | - Sex | -0.114 | 0.472 |
| | - Age | -0.169 | 0.285 |
| | - Body mass index | 0.241 | 0.125 |
| | - Arm calibration order | 0.076 | 0.633 |
| | - First block conditioning | -0.038 | 0.811 |
| | - Last block conditioning | -0.152 | 0.336 |
| | - Test stimulus pressure | 0.381 | 0.013 * |
| | - # calibration stimuli test pressure | -0.117 | 0.479 |
| | - Test stimulus calibration slope | -0.376 | 0.018 * |
| | - Conditioning stimulus peak pressure | 0.242 | 0.123 |
| | - Conditioning stimulus trough pressure | 0.229 | 0.145 |
| | - # calibration stimuli conditioning stimulus | 0.018 | 0.914 |
| | - Conditioning stimulus calibration slope | -0.230 | 0.158 |

Rated CPM: higher values reflect more hypoalgesia. Verbal CPM: 1 = hypoalgesia, 0 = no difference, -1 = hyperalgesia. * $p < .05$, no correction for multiple comparisons.
